# LigEGFR: Spatial graph embedding and molecular descriptors assisted bioactivity prediction of ligand molecules for epidermal growth factor receptor on a cell line-based dataset

**DOI:** 10.1101/2020.12.24.423424

**Authors:** Puri Virakarin, Natthakan Saengnil, Bundit Boonyarit, Jiramet Kinchagawat, Rattasat Laotaew, Treephop Saeteng, Thanasan Nilsu, Naravut Suvannang, Thanyada Rungrotmongkol, Sarana Nutanong

**Affiliations:** Kamnoetvidya Science Academy (KVIS), Rayong 21210, Thailand; School of Information Science and Technology, Vidyasirimedhi Institute of Science and Technology (VISTEC), Rayong 21210, Thailand; Program in Bioinformatics and Computational Biology, Graduate School, Chulalongkorn University, Bangkok 10330, Thailand; Biocatalyst and Environmental Biotechnology Research Unit, Department of Biochemistry, Faculty of Science, Chulalongkorn University, Bangkok 10330, Thailand

**Keywords:** Deep learning, Machine learning, Graph embedding, Drug discovery, QSAR, Epidermal growth factor receptor

## Abstract

**Motivation:** Lung cancer is a chronic non-communicable disease and is the cancer with the world’s highest incidence in the 21^st^ century. One of the leading mechanisms underlying the development of lung cancer in nonsmokers is an amplification of the epidermal growth factor receptor (EGFR) gene. However, laboratories employing conventional processes of drug discovery and development for such targets encounter several pain-points that are cost- and time-consuming. Moreover, high failure rates are caused by efficacy and safety problems during research and development. Therefore, it is imperative to develop improved methods for drug discovery. Herein, we developed a deep learning model with spatial graph embedding and molecular descriptors based on predicting pIC_50_ potency estimates of small molecules and classifying hit compounds against the human epidermal growth factor receptor (LigEGFR). The model was generated with a large-scale cell line-based dataset containing broad lists of chemical features.

**Results:** LigEGFR outperformed baseline machine learning models for predicting pIC_50_. Our model was notable for higher performance in hit compound classification, compared to molecular docking and machine learning approaches. The proposed predictive model provides a powerful strategy that potentially helps researchers overcome major challenges in drug discovery and development processes, leading to a reduction of failure to discover novel hit compounds.

**Availability:** We provide an online prediction platform and the source code that are freely available at https://ligegfr.vistec.ist, and https://github.com/scads-biochem/LigEGFR, respectively.

**Key points:**

- LigEGFR is a regression model for predicting pIC_50_ that was developed for the human EGFR target. It can also be applied to hit compound classification (pIC_50_ ≥ 6) and has a higher performance than baseline machine learning algorithms and molecular docking approaches.
- Our spatial graph embedding and molecular descriptors based approach notably exhibited a high performance in predicting pIC_50_ of small molecules against human EGFR.
- Non-hashed and hashed molecular descriptors were revealed to have the highest predictive performance by using in a convolutional layers and a fully connected layers, respectively.
- Our model used a large-scale and non-redundant dataset to enhance the diversity of the small molecules. The model showed robustness and reliability, which was evaluated by y-randomization and applicability domain analysis (ADAN), respectively.
- We developed a user-friendly online platform to predict pIC_50_ of small molecules and classify the hit compounds for the drug discovery process of the EGFR target.

## 1 Introduction

Non-communicable diseases (NCDs) are increasingly leading causes of death. Lung cancer is a global health burden and has been reported as the most frequent cancer worldwide; in 2018, lung cancer was the most commonly diagnosed cancer (11.6% of the total cases) and the leading cause of cancer death (18.4% of the total cancer deaths) [1]. The leading causes of lung cancer in nonsmokers and those who are not in close contact with smokers are the amplification and hyperactivation of the epidermal growth factor receptor (EGFR). EGFR is a member of receptor tyrosine kinases (RTKs) family located on the cell membrane and the elevated activity of this enzyme results in increased cascades of the RAS-RAF-MEK-ERK and PI3K-AKT-mTOR signaling pathways [2]. Chemotherapy is the traditional treatment, with drugs taken at high doses, but it can cause a number of unpleasant side effects such as dizziness, alopecia, oral and pharyngeal tissue damage, and fatigue [3]. Nowadays, targeted therapy is a new approach to treatment that is developed specifically for cancer targets and has generated agents such as Gefitinib, Erlotinib, Lapatinib, Afatinib, Dacomitinib, Neratinib, Osimertinib, and Brigatinib [4].

Computer-aided drug discovery can help reduce the time and cost in the early stages of drug discovery and development process [5]. The virtual screening technique is a method for accelerating drug discovery and is commonly performed by molecular docking, pharmacophore modeling and mapping, and molecular similarity calculations. However, these traditional techniques have several disadvantages, such as the inability to identify whether the compound is an inhibitor or an activator for the target. The ligand with the highest score from molecular docking does not automatically have the highest potential to be a useful lead compound [6]. Over the past decade of drug discovery, the use of big data sets, such as bioactivity and drug-target interactions, has been dramatically increased through the development of high-throughput screening (HTS) technologies [7, 8, 9]. Consequently, machine learning and deep learning techniques now play important roles in the pharmaceutical industry by enhancing efficiency in hit compound discovery with a result in decreasing the proportion of drug failures in clinical trials.

Machine learning is a widely used tool that opens vast opportunities in the drug development process. It can be applied to constructing QSAR models, built through various state-of-the-art techniques. QSAR explores the structure of the molecule to predict the activities, such as bioactivity (IC_50_, EC_50_, K_i_, and K_d_), adsorption, distribution, metabolism, elimination, and toxicity (ADMET) properties; it is an essential indicator in selecting lead compounds before they are tested *in vitro* and *in vivo*. Generally, QSAR studies in drug discovery are focused on the physicochemical properties of the molecules that trigger similar biological effects [10]. There are prominent published works that used deep learning for predicting molecular properties such as binding affinity [11, 12, 13], drug-target interaction [14], and bioactivity [15]. It can be seen that bioactivity prediction using machine learning methods is helpful in accelerating the drug discovery process. For EGFR, there have been various machine learning models developed to predict IC_50_ of ligands against wild type EGFR that were based on small scale datasets (up to 290 compounds) [16, 17, 18, 19, 20, 21, 22]. But machine learning models that are based on small-scale compound datasets may not accurately predict properties owing to out-of-feature domains and low diversity of substructure features in the dataset molecules.

To tackle the problems and pitfalls in the previous methods of drug discovery, we first developed a machine learning model to predict pIC_50_ of small molecules using the human EGFR as the target and termed it LigEGFR. The development of this model using a large-scale cell line-based and non-redundant dataset is described here. The convolution spatial graph embedding network (C-SGEN) [23] [24] together with the deep neural network (DNN) technique was modified and applied for model training in this work. C-SGEN is comprised of convolution spatial graph embedding layers (C-SGELs) which are constructed by graph convolutional networks. C-SGEL is introduced to maintain the spatial relationship of atoms in a molecule by using the GraphCov features [25]. The molecular graph represented by an adjacency matrix and a node matrix is used as an input of C-SGELs.

The main contributions of our work are as follows:

i. We employed the C-SGEN algorithm with architecture adaption and DNN for pIC_50_ prediction with a large-scale and non-redundant dataset of bioactive compounds against human EGFR.
ii. The non-hashed and hashed molecular fingerprints were considered for convolutional (Conv) layers and fully connected (FC) layers, respectively.
iii. We constructed a user-friendly web service with compatibility for all devices, https://ligegfr.vistec.ist, and Python executable script, https://github.com/scads-biochem/LigEGFR, for predicting pIC_50_ and classifying hit compounds of small molecules against human EGFR.

## 2 Methods

The overview of LigEGFR is shown in Figure 1. We used the large cell line-based and non-redundant dataset to retrieve broad features of small molecules. These features were applied to LigEGFR, comprising C-SGELs for the GraphCov features, FC layers, and Conv layers for different fingerprint combinations. LigEGFR was then compared to the baseline machine learning algorithms: C-SGEN, CNN, and random forest (RF). Also, the classification performance of the conventional virtual screening techniques such as molecular docking (CCDC GOLD and AutoDock Vina programs), was determined. The model assessment was performed by the y-scrambling technique and applicability domain analysis (ADAN). The details of the methodologies are particularized in the following subsections. For web service construction, the description is provided in the supplemental section S1^†^.

**Figure 1:**
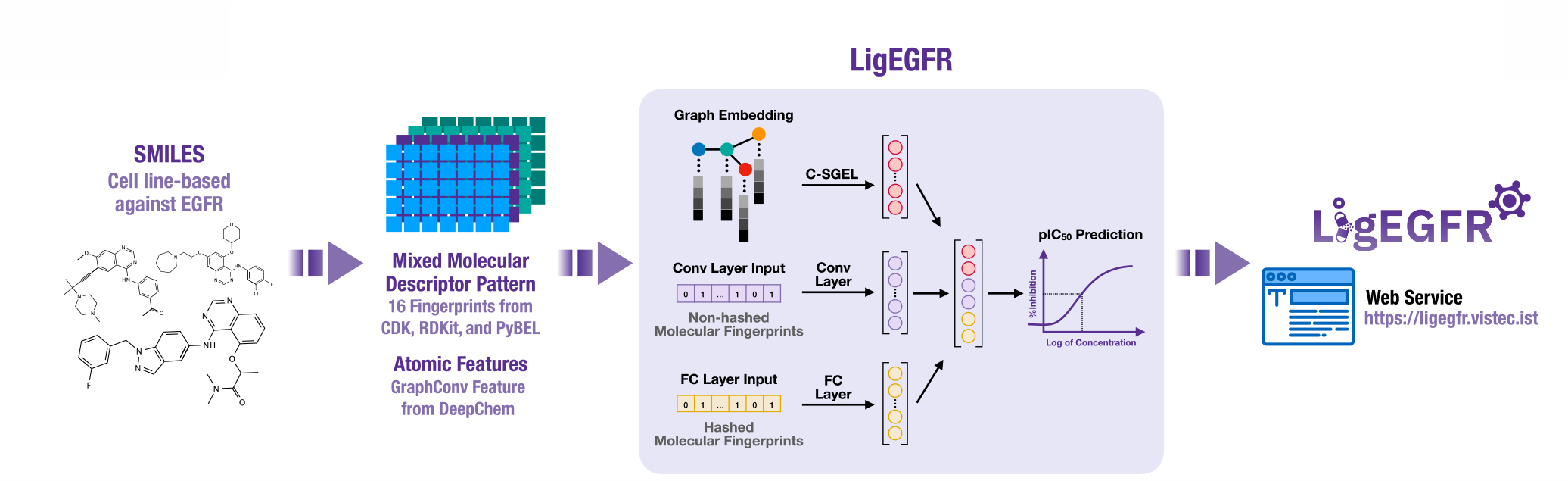
This overview represents LigEGFR construction. The dataset, SMILES strings, is converted into the GraphCov features, non-hashed molecular fingerprints, and hashed molecular fingerprints. LigEGFR encodes these features to construct a predictive model by using C-SGEL for the GraphCov features, the Conv layers for the non-hashed molecular descriptors, and the FC layers for the hashed molecular descriptors.

### 2.1 Data preparation

The 37,753 substances (90,118 data points) known to inhibit the activity of human EGFR were collected from the Reaxys database [26]. The crucial information consisted of the ligand structures in the format of SMILES (Simplified Molecular-Input Line-Entry System) and the pIC_50_ values derived as −*log*(IC_50_). The data from wild type human EGFR and *in vitro* experiment as well as from cell-based assay were selected. The redundant compounds with the same SMILES and those with missing pIC_50_ values or SMILES were removed; salt derivatives in SMILES was also eliminated. After that, we removed outliers by considering the rule of five (RO5) criterion, with (i) hydrogen bond donors, (ii) hydrogen bond acceptors, (iii) molecular weight, and (iv) the octanol-water partition coefficient (*log* P). We used the interquartile range method (IQR) by

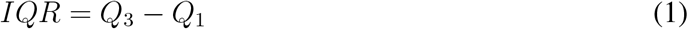

and outliers were qualified for exclusion if they fell outside the range as follows:

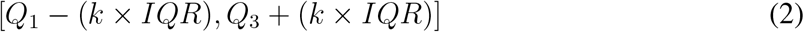

where *k* = 3. In this process, Pandas [27], NumPy [28], Scipy [29], Matplotlib [30], scikit-learn [31], RDKit [32], PyBEL [33], and Seaborn [34] packages were used for data management and processing.

### 2.2 Molecular fingerprints

The SMILES of compounds were encoded into a molecular fingerprint comprising a set of molecular descriptors or features. In this work, we made 16 molecular fingerprints from the CDK [35], RDKit [32], and PyBEL [33] packages (see supplemental Table S1^†^). These descriptors were represented in Boolean form. The types of molecular fingerprints were divided into two groups, non-hashed and hashed molecular fingerprints. Furthermore, the fingerprints were reduced in dimension with different correlation coefficients in feature selection and the number of principal components in the PCA method. For the feature selection method, the variance threshold was fixed to 0.001, and the correlation coefficient was varied from 0.75 to 0.90. For the PCA method, the number of components was altered from 50 to 800.

### 2.3 Dataset splitting

Our dataset was divided into training set and test set using the Train/Test split method [31]. The training set ratio to the test set was 80:20, where 80% of the final dataset was used as an internal set (2,794 compounds) and the remaining 20% used for the test set were unseen data (699 compounds). The internal set was then separated into a training set (72%, 2,515 compounds) and validation set (8%, 279 compounds).

### 2.4 Model development

Figure 2 shows the structure of a LigEGFR architecture. The C-SGEN algorithm [23] was applied for LigEGFR construction with a modification in the architecture. Normally, each molecule is represented by a molecular graph and hashed and non-hashed molecular descriptors. The connection between atoms is constructed as a graph that is initialized from the node matrix (*X*) and an adjacency matrix (*A*). By regarding the atomic connection between atoms *i* and *j*, if it appears then *A*_*i,j*_ = 1; otherwise, *A*_*i,j*_ = 0. The initial node matrix (*X*) is proposed by a set of atoms *x*_*i*_:

**Figure 2:**
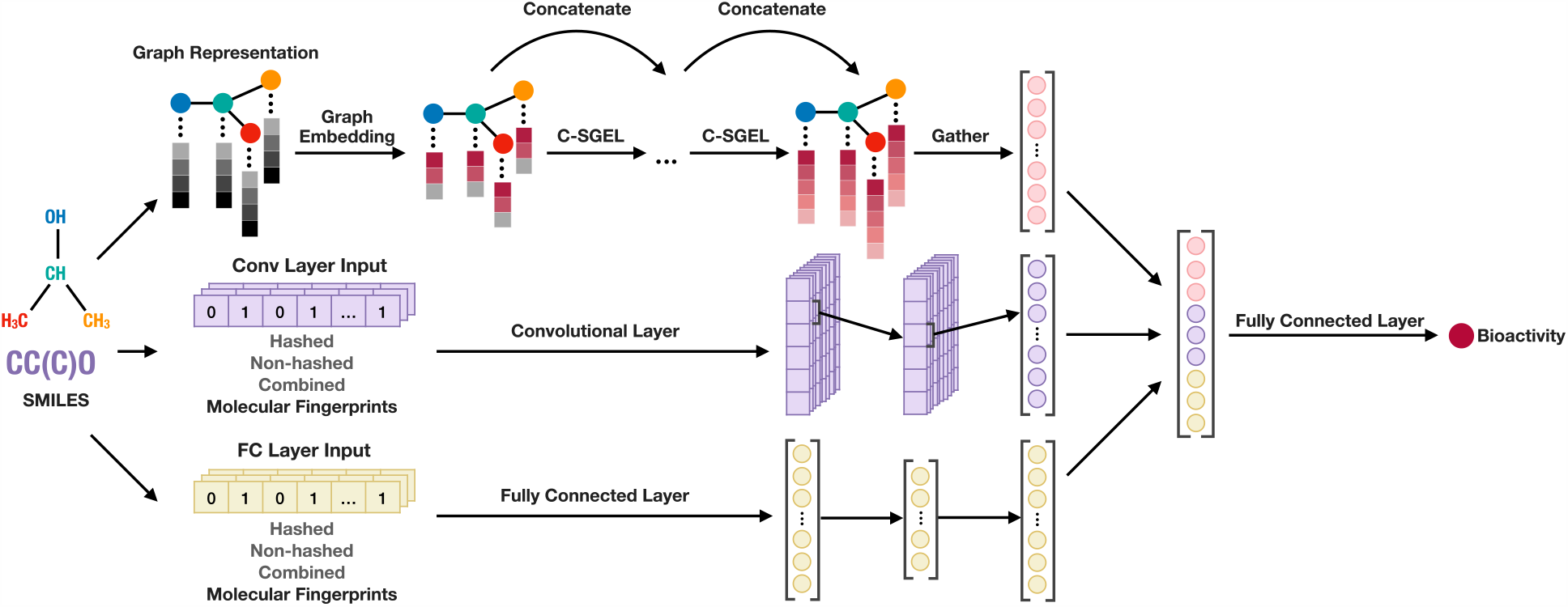
Diagram of the LigEGFR architecture contains the C-SGEN algorithm with architecture adaptation for graph embedding and the DNN algorithm for molecular fingerprints.

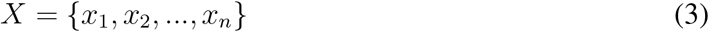

where each atom *x*_*i*_ is an *m*-dimensional vector. The vector is extracted from DeepChem [25] and encoded into a one-hot vector by the atomic features shown in supplemental Table S2^†^. The propagation function used in C-SGEN follows the equation

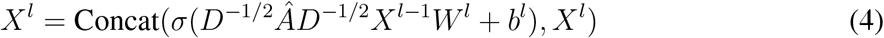

where *Â* = *A* + *I*_*n*_ and *D* is the diagonal matrix containing degree of each node in *Â. D*^−1/2^*ÂD*^−1/2^ is a symmetric normalization of an adjacency matrix, where the propagation of each adjacent node (*i, j*) is normalized by the degree of both nodes *i* and *j* [36]. 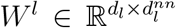 and 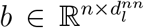 represent the learnable weight and bias, respectively, where *d*_*l*−1_ is the dimension of layer *l* − 1 and 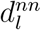 is the dimension resulted from the neural network computation in layer *l*. The feature vector resulted from the neural network is a matrix with dimension 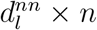 concatenated to the original feature vector *X*^*l*−1^ resulted in the output matrix *X*^*l*^ with dimension 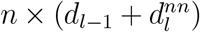. The illustration of C-SGEN propagation is shown in Figure 3.

**Figure 3:**
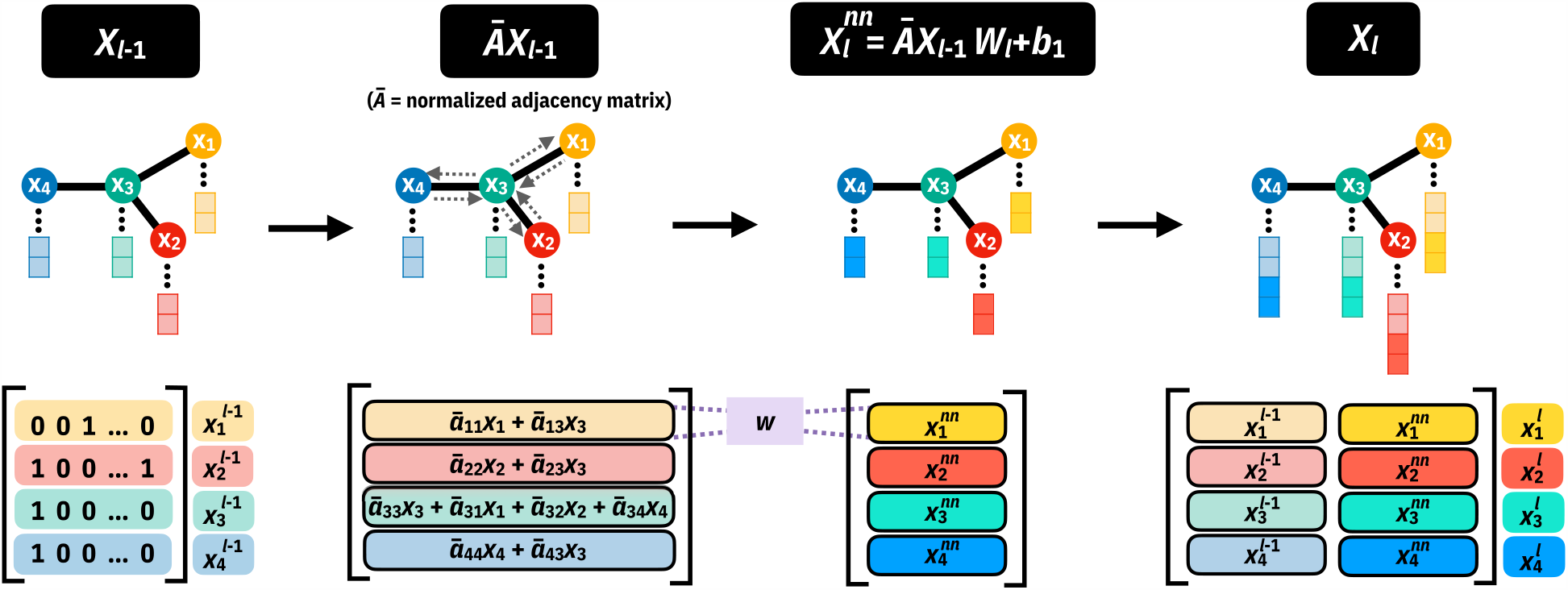
This schematic of C-SGEN propagation contains a visualization of the step in graph feature propagation that includes the multiplication of the feature with the normalized adjacency matrix *Ā* which is equal to *D*^−1/2^*ÂD*^−1/2^. Afterward, the hidden learnable computation layer of weight and bias is used to extract the feature and concatenate back to the previous layer.

After C-SGEN, the feature matrix *X*^*f*^ comprises each atom feature vector 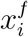, where *f* is the final layer of C-SGEN. In order to obtain an order-invariant feature for one molecule, the molecular graph representation *x*_*graph*_ is calculated by a summation of all atomic features 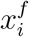 as follows.

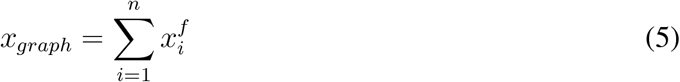

Here the dimension of molecular graph representation *x*_*graph*_ is the same as the atomic feature vector 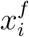. Afterward, the molecular fingerprint is used as another input of the model to confirm the feature’s generalization. These molecular fingerprints can be split into hashed and non-hashed fingerprints for the input of the Conv layers and FC layers. The fingerprints of each molecule can be expressed as *x*^0^, and the output from the *l*th hidden layer can be represented as *x*^*l*^ using the equation.

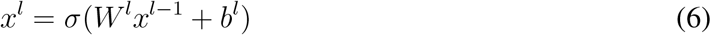

where *σ, W* ^*l*^ and *b*^*l*^ are activation function, learnable weight matrix, and bias for the *l*th hidden layers, respectively. Then, the output from the last layer of both Conv and FC layers are concatenated together into an updated molecular fingerprint, denoted by *x*_*finger*_.

Lastly, *x*_*graph*_ and *x*_*finger*_ are concatenated into the final molecular representation, *x*_*mol*_. This *x*_*mol*_ is used to predict the pIC_50_ (*y*_*predicted*_) of the molecule through the FC layers as follows.

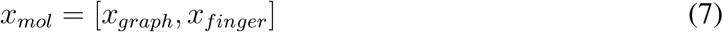

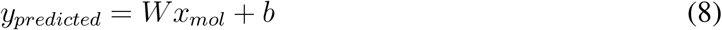

where *W* is the weight matrix and *b* is the bias.

To find the fingerprint pattern that is followed through with a high-performance model, we have changed the combination of fingerprints used as input to the Conv layers and FC layers as shown below:

1. The hashed and non-hashed fingerprints were applied in the Conv layers and FC layers, respectively.
2. The hashed and non-hashed fingerprints were used in the FC layer and Conv layers, respectively.
3. Both hashed and non-hashed fingerprints (all fingerprints) were applied in both FC and Conv layers.

In addition, hyperparameter tuning was applied to this architecture by adjusting *BatchSize, lr, ch_num, k*, and *csgellayer*. The detail of hyperparameter tuning is described in supplemental section S2^†^.

### 2.5 Model assessment

#### 2.5.1 Predictive performance

We evaluated the performance of the regression models by using the coefficient of determination (*R*^*2*^) and root mean square error (*RMSE*).

The *R*^*2*^ indicates how well the predicted values compared with the observed values

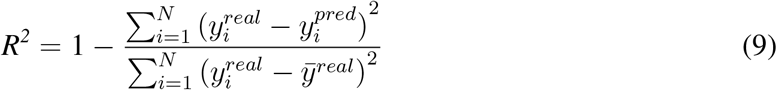

while the *RMSE* represents the root mean square or quadratic mean of the difference between the predicted value and the observed value.

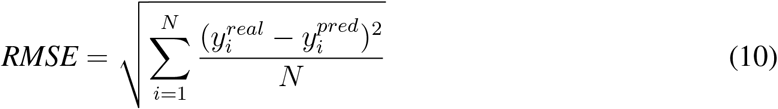

where *y*^*real*^ is the observed pIC_50_ value from experiments, *y*^*pred*^ is the predicted pIC_50_ value from the machine learning model, and *N* is the number of compounds used for the performance test. Each metric was used to evaluate the training set (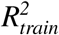 and *RMSE*_*train*_), validation set (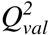 and *RMSE*_*val*_) and test set (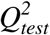 and *RMSE*_*test*_) by averaging statistical values and changing the random seed.

#### 2.5.2 Y-scrambling validation

Basically, the robustness of the model is evaluated by the y-scrambling or y-randomization technique. This procedure can predict the performance of the model by comparing the observed values (*y*^*real*^) to the model built from shuffled *Y* – leaving the entire *X* [37, 38]. If the original descriptors cannot predict the random pIC_50_ value which is calculated from the shuffled model, the original model is robust. The basic leave-one-out (LOO) technique was applied for statistics of the scrambled models. In this experiment, we employed the 100 y-scrambling models for comparing 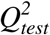 to the initial model.

#### 2.5.3 Applicability domain analysis (ADAN)

The applicability domain (AD) is the chemical space, structure, or knowledge from a training set in which the model makes reliable predictions and is considered to be applicable [39]. The AD determines the extent of the data for which the predictions of the model are valid. In other words, the prediction must be an interpolation. Typically, the AD is calculated by a range-based method, geometric method, distance-based method, or probability density distribution-based method [40]. We used k-nearest neighbors (kNN)-based and PCA bounding box methods for evaluation. These methods are simple to understand and interpret. In the kNN-based method, if the Euclidean distances between a query and the training set exceed the ADT, the prediction is not reliable, and *vice versa*. ADT was calculated as follows for new substance prediction [41].

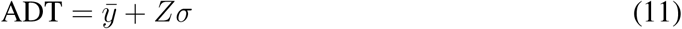

Here, 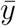 is an average of the Euclidean distances of kNN in the training set, in which *k* = 50 as the square root of the training set number [42], *Z* is an arbitrary parameter (0.5 by default), and *σ* is a standard deviation of the Euclidean distances.

For the PCA bounding box, a 3D principal component (PC) space was generated for the training set, test set, approved drug set, and clinical trial compound set. Each point in the PC space represents a compound, so the training set’s chemical space can be found. If a query is in the bounding box, the prediction is reliable (interpolation) [43].

### 2.6 Baseline comparison

We have compared the efficiency of our LigEGFR with baseline machine learning algorithms and the conventional molecular docking methods in predicting pIC_50_ value and whether a query is a hit compound (pIC_50_ ≥ 6) [44]. The receiver operating characteristic (ROC) curves, plotting the true-positive rate against the false-positive rate, were obtained from both approaches by changing the threshold used to classify each ligand [45]. Moreover, the enrichment plot, plotting the percentage of identified hit compounds against the selected top compounds, indicates how rich the correctly predicted hit compound is in the whole ranked dataset. The area under the ROC curve (AUC), and the enrichment plot were used as metrics for performance comparison. Furthermore, the balanced-accuracy, precision, recall, F1-score, and Matthews correlation coefficient (MCC) were also evaluated to determine the efficiency of the classification task. The definitions of these metrics are shown below.

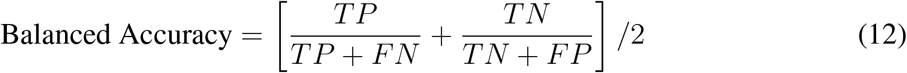

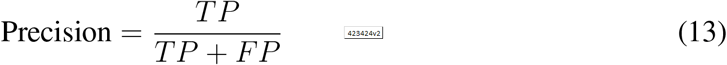

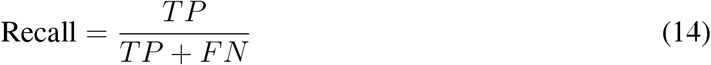

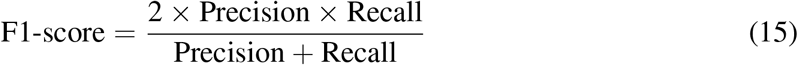

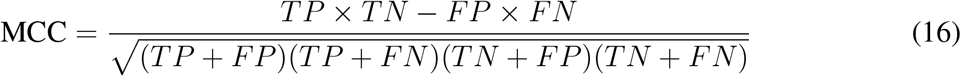

*TP* (true-positives) and *FN* (false negatives) denote the numbers of known active compounds predicted to be active compounds (pIC_50_ ≥ 6) and inactive compounds (pIC_50_ < 6). *TN* (true negatives) and *FP* (false positives) indicate the numbers of known inactive compounds predicted to be inactive compounds and active compounds.

#### 2.6.1 Machine learning-based approaches

The baseline evaluation was determined from machine learning techniques by RF, CNN, and C-SGEN algorithms. We developed the RF model by using the recursive feature elimination (RFE) technique with *Adjusted R*^*2*^. The CNN model was applied using Conv and FC layers with reduced hashed and non-hashed fingerprints. The C-SGEN algorithm was performed as a controlled experiment.

Additionally, RF and CNN models were tuned by hyperparameter optimization. The detailed methods are described in supplemental section S3^†^, S4^†^, and S5^†^ for RF, CNN, and C-SGEN algorithms, respectively.

#### 2.6.1 Molecular docking-based approaches

We applied the molecular docking to calculate fitness scores between a list of test set ligands and EGFR protein. Generally, the ligand with the lowest binding free energy (ΔG_bind_) which has the highest binding affinity to the drug target has the greatest chance to be an efficient hit compound. Each ligand’s fitness score from the test set was obtained from the EGFR tyrosine kinase in both active and inactive conformations: PDB entry codes 1M17 [46] and 1XKK [47]. These protein structures were prepared using the python script, *clean_pdb_keep_ligand*.*py* through Rosetta3 [48]. All ligand geometries in the test set were optimized by the MMFF94 force field with RDKit and subsequently converted into the .mol2 file. The CCDC GOLD (the Genetic Optimisation for Ligand Docking) version 5.8.0 [49] with ChemPLP (Piecewise Linear Potential) fitness score [50], and AutoDock Vina 1.1.2 [51] were implemented for molecular docking. The described details of molecular docking are provided in supplemental section S6^†^ and S7^†^.

## 3 Results and discussion

### 3.1 Predictive performance for regression tasks

As shown in Table 1, the LigEGFR model, after tuning the hyperparameter with the feature selection technique, outperformed all baseline algorithms for the regression task (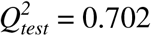 and *RMSE* = 0.802). The best hyperparameters for LigEGFR were *BatchSize* = 32, *lr* = 5 × 10^−5^, *ch_num* = 4, *k* = 8, and *csgellayer* = 4. More importantly, LigEGFR also provided a more accurate pIC_50_ prediction for the test set compared to those of the baseline algorithms (see Figure 4). The predictive performance of the test set described by 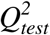 in QSAR task, should be ≥ 0.6 [52]. LigEGFR composes graph embedding layers that are gathered from an atomic feature representation of the molecule, which is based on the C-SGEN algorithm [23]. The advantage of the graph-based method for molecular representation is that it is able to pull out essential features of molecular information including the neighboring atoms.

**Table 1:**
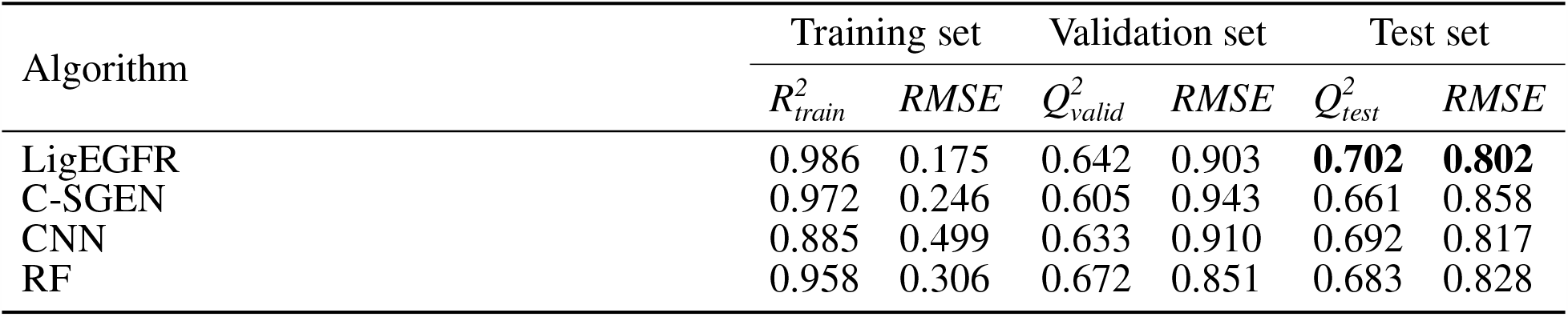
LigEGFR, C-SGEN, CNN, and RF models’ predictive performance is indicated for regression tasks. The results of all models were computed and tuned with the various hyperparameters, with the exception of C-SGEN which was trained on the original hyperparameter.

**Figure 4:**
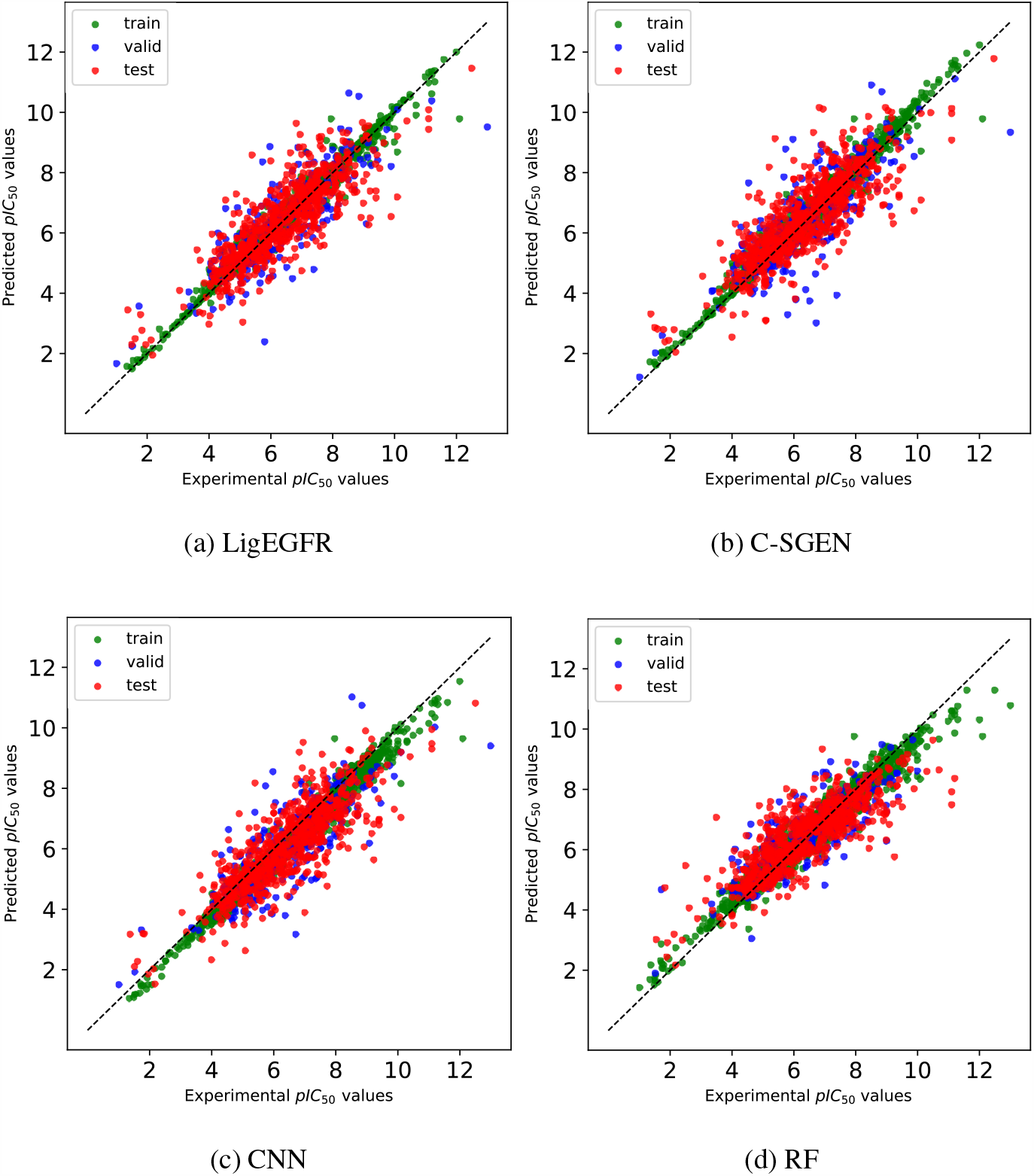
Comparison of relationships between predicted and experimental pIC_50_ values generated by for (a) LigEGFR, (b) C-SGEN, (c) CNN, and (d) RF. The dashed line shows where the points should ideally lie.

The additive non-hashed and hashed molecular descriptors embedded on Conv and FC layers, have improved the predictive performance and generalized model. The feature analysis of these molecular descriptors is described in supplemental section S8^†^. From Table 2, we noticed that non-hashed molecular descriptors on the Conv layers outperformed other arrangements in predictive performance. The non-hashed molecular descriptor is a feature that represents the presence of the substructures of a molecule such as atom type, neighboring atoms, atomic groups, bonds, etc. The Conv layers with substructure molecular fingerprints may learn relationships and similar features of the various small molecules to attain a global representation of the information that influences the physicochemical properties of the molecule. The non-hashed molecular fingerprints on the Conv layers and the hashed molecular fingerprints on the FC layers with cut-off values by feature selection, 0.001 for low variance and 0.90 for collinearity, exhibit the highest predictive performance (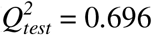 and *RMSE*_*test*_ = 0.813) without hyperparameter tuning.

**Table 2:** The predictive performance based on LigEGFR, trained by Conv and FC layers with a combination of molecular fingerprint patterns. The feature selection technique was completed for molecular fingerprints prior to hyperparameter optimization.

| Pattern of molecular fingerprint |  |  | Predictive performance |  |  |  |  |  |
| --- | --- | --- | --- | --- | --- | --- | --- | --- |
|  |  |  | Training set |  | Validation set |  | Test set |  |
| Conv layers | FC layers | Cut-off value | $R^2_{train}$ | $RMSE$ | $Q^2_{valid}$ | $RMSE$ | $Q^2_{test}$ | $RMSE$ |
| Hashed | Non-hashed | [0.001, 0.75] | 0.996 | 0.099 | 0.617 | 0.929 | 0.649 | 0.873 |
| Non-hashed | Hashed | [0.001, 0.75] | 0.994 | 0.117 | 0.642 | 0.899 | 0.693 | 0.816 |
| All | All | [0.001, 0.75] | 0.995 | 0.109 | 0.632 | 0.911 | 0.689 | 0.822 |
| Hashed | Non-hashed | [0.001, 0.80] | 0.996 | 0.099 | 0.623 | 0.922 | 0.656 | 0.865 |
| Non-hashed | Hashed | [0.001, 0.80] | 0.994 | 0.120 | 0.639 | 0.902 | 0.694 | 0.815 |
| All | All | [0.001, 0.80] | 0.995 | 0.108 | 0.628 | 0.916 | 0.691 | 0.818 |
| Hashed | Non-hashed | [0.001, 0.85] | 0.996 | 0.100 | 0.628 | 0.916 | 0.659 | 0.860 |
| Non-hashed | Hashed | [0.001, 0.85] | 0.993 | 0.125 | 0.640 | 0.902 | 0.692 | 0.819 |
| All | All | [0.001, 0.85] | 0.994 | 0.112 | 0.628 | 0.916 | 0.690 | 0.821 |
| Hashed | Non-hashed | [0.001, 0.90] | 0.996 | 0.100 | 0.627 | 0.917 | 0.663 | 0.855 |
| Non-hashed | Hashed | [0.001, 0.90] | 0.993 | 0.125 | 0.636 | 0.906 | **0.696** | **0.813** |
| All | All | [0.001, 0.90] | 0.994 | 0.112 | 0.624 | 0.920 | 0.694 | 0.814 |

Note: Cut-off values [I,II]; I: low variance of molecular descriptors less than defined value will not be considered. II: collinearity of molecular descriptors more than a defined value will not be considered.

In comparing the C-SGEN algorithm with the original hyperparameter, this algorithm has influences that decrease the predictive performance because the dataset did not conform to the original hyperparameter setting. Moreover, the additive non-hashed and hashed molecular descriptors were not applied to Conv and FC layers, thus it was inadequate for predicting the pIC_50_ value. The types and shapes of features are prominent requirements for a learning process using proper hyperparameters and machine learning algorithms. The CNN model with only molecular descriptors can predict pIC_50_, because this model was optimized with the hyperparameters and embedded with non-hashed and hashed molecular descriptors into different FC and Conv layers. For the RF model, molecular descriptors have been shown to have high performance in predicting pIC_50_, quite equivalent to the CNN algorithm, because a molecular descriptor as a binary digit has the ability to participate in the ensemble learning process on the decision tree [53]. Consequently, the molecular graph information on C-SGEL plays a significant role in enhancing the predictive performance of the regression model.

All results of hyperparameter tuning for LigEGFR, CNN, and RF algorithms are available in supplemental Table S7^†^, S8^†^, and S9^†^, respectively.

### 3.2 Reliability validation of models

#### 3.2.1 Y-scrambling

The y-scrambling is a statistical technique for assessing the robustness of the models. Wold *et al*. [54] described a criterion to decide an agreed model, based on the 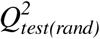 and 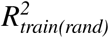. If the test provides 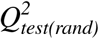 and 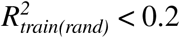, a result in the model is not a chance correlation to predict a scrambled *Y* value. Figure 5 shows the plots that the models obtained from actual data (blue dot) arranged in the top right quadrant, thereby suggesting robust models. Meanwhile, the red dots are in the bottom left quadrant (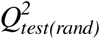 and 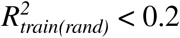), indicating the low-performance to predict shuffled *Y* values calculated from the original *X*. Hence, our LigEGFR and other models trained on the real dataset have high robustness.

**Figure 5:**
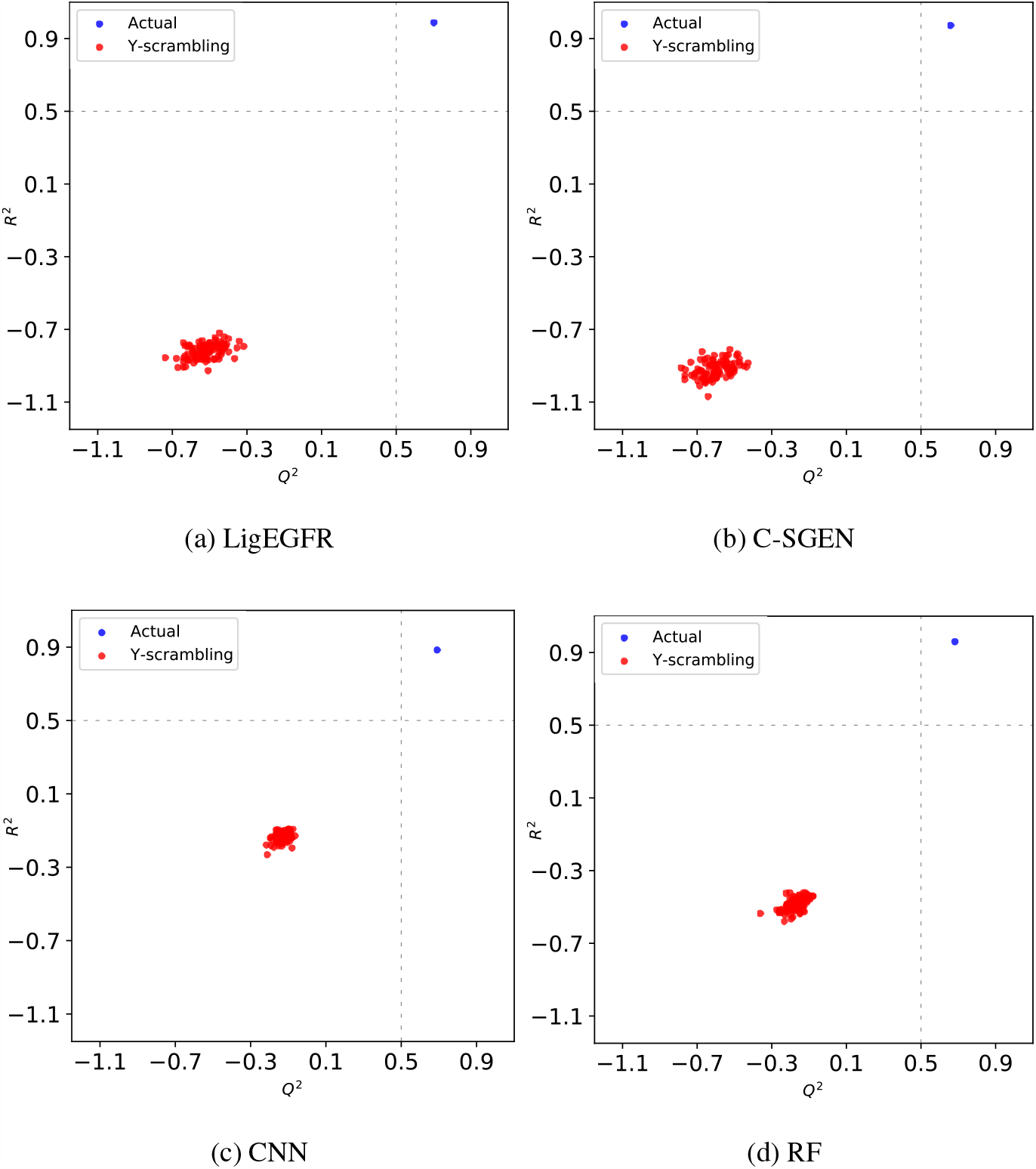
Scatter plots of y-scrambling validation for (a) LigEGFR, (b) C-SGEN, (c) CNN, and (d) RF models.

#### 3.2.2 Loss curves

Generally, the correctness of the error function of neural networks is indicated by the loss function, which must be minimized. The loss curve emerges to explain model learning and model performance by optimum, overfitting, or underfitting. In this experiment, *MSE* or *L2* was used for the loss function. The *MSE* is mostly applied for regression problems, estimated by the average of squared differences between the predicted and real values. As shown in Figure 6, the loss curves presented a sharp drop to reach an optimal value in the training set. Meanwhile, the loss curves of the validation set remained constant, a convergence state. We point out that all models were not overfitting, since the loss curves of the validation set and training set were consistent.

**Figure 6:**
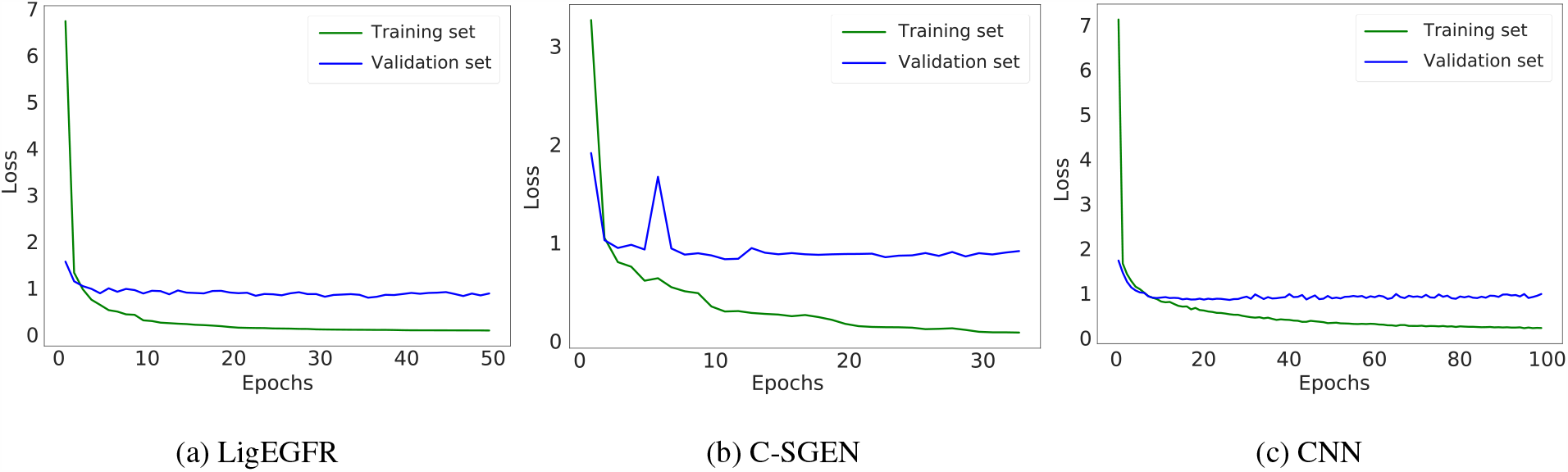
Loss curves of the training set and validation set are indicated by the green and blue curves, respectively.

#### 3.2.3 Applicability domain analysis (ADAN)

Evaluating the model’s applicability, *i*.*e*. assessing whether it is appropriate to the diversity of molecular scaffolds and functional groups, is a crucial step in QSAR method development. We note the results of ADAN by the PCA bounding box and the kNN-based methods, in Figure 7 and Table 3, respectively. The kNN-based method provides quantitative data of AD; the PCA bounding box method gives visual and qualitative information of AD. For the kNN-based method, more than 70% of the data in the test set, approved drug set, and clinical trial compound set were in the applicability domain. For the PCA bounding box method, it can be clearly seen that the majority of the data in all sets are in the domain of the training set or interpolation. The results showed that the training set has a large chemical space representing the test set, commercial drugs, and clinical trial drugs. Consequently, our model can reliably predict a new substance as having a high chance to be a lead compound.

**Table 3:** Applicability domain analysis with the kNN-based method for the test set, approved drug set, and clinical trial compound set, where *k* = 50

|  | Test set | Approved drug set | Clinical trial compound set |
| --- | --- | --- | --- |
| Number of compounds | 699 | 8 | 18 |
| Reliability | 72.82% | 87.50% | 72.22% |

**Figure 7:**
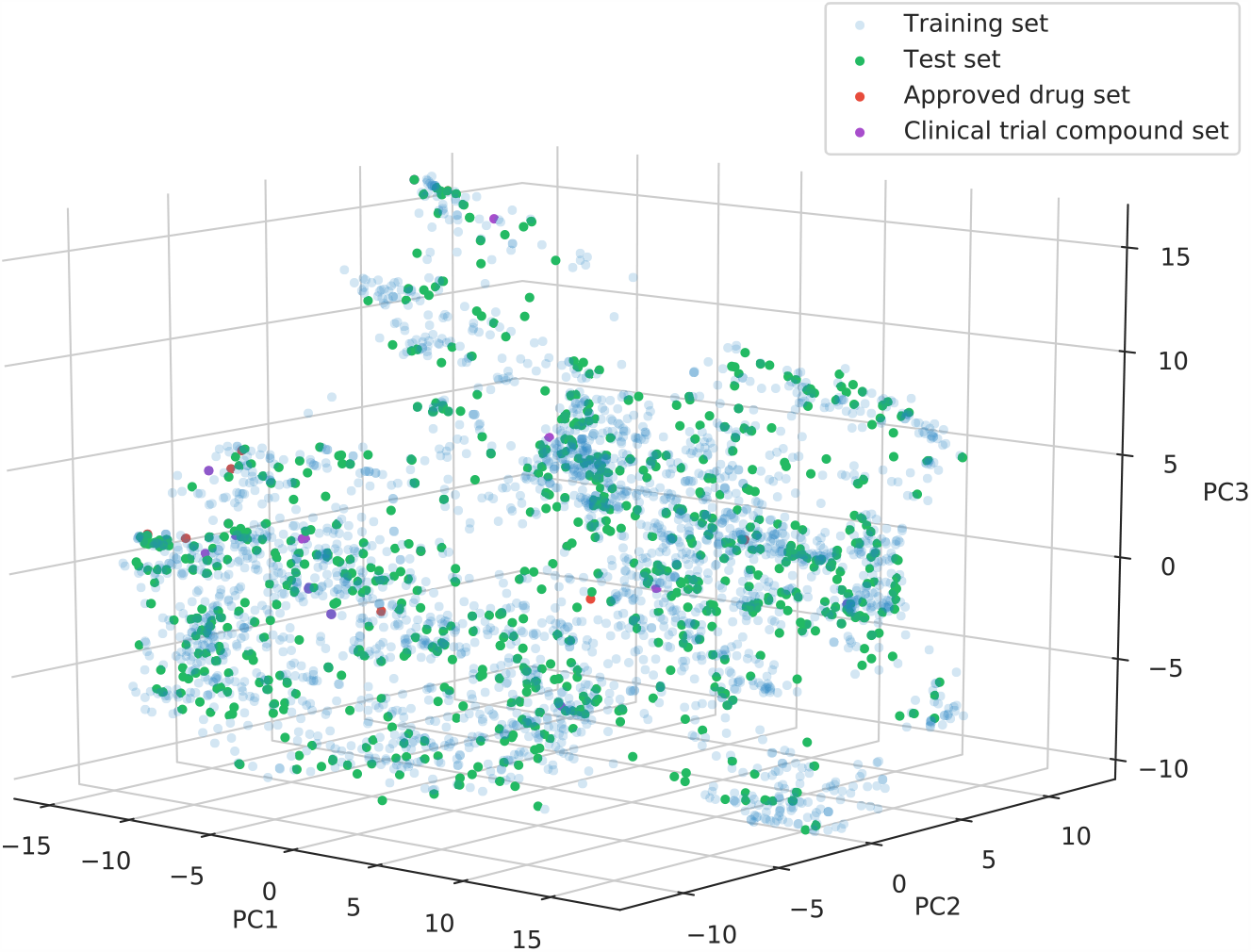
The 3D visualization of the applicability domain shown by the PCA bounding box method.

### 3.3 Predictive performance for classification tasks

The utility of virtual screening can be compared to molecular docking in the task of classifying active and inactive compounds. Wang *et al*. [55] and Plewczynski *et al*. [56] have recognized that the CCDC GOLD has the best accuracy for the top-scored pose, while Autodock Vina has the best scoring power for both the top-scored and the best pose.

As listed in Table 4, the performance in the classification of hit compounds (pIC_50_ ≥ 6) suggested that LigEGFR performs with the highest balanced accuracy (0.867) and MCC (0.732) compared to the baseline machine learning algorithms. The balanced accuracy is the number of correct predictions divided by the total number of compounds in an imbalanced dataset. Normally, overall accuracy is not a suitable metric for a dataset that is imbalanced [57]. In our test set, all ligands contained 419 active and 280 inactive compounds. MCC is a useful metric to measure performance for binary classification problems. Chicco and Jurman [58] suggested that the MCC works as a truthful metric for the evaluation of binary classification rather than for accuracy and F1-score for imbalanced datasets. Emphasis on overall accuracy and F1-score can be a prevalent mindset leading to mis-interpretation of results, due to lack of consideration of the ratio between positive and negative constituents. In Figure 8a, the ROC plot indicates the classification performance on a different threshold. The result shows that LigEGFR provides AUC at 0.942, which is slightly better than the baseline methods in machine learning (0.917 – 0.934) and significantly higher than molecular docking techniques (0.466 – 0.628). Additionally, enrichment plots were used for evaluating how a model can perform in ranking pIC_50_ value. The enrichment plot was employed by considering the top-ranked data and finding how many active compounds there are. The result showed that the true active compounds, identified by percentages in the LigEGFR model, were mostly higher than others at the same top-ranked dataset percentages (see Figure 8b), which was in an accordance with the ROC plots.

**Table 4:**
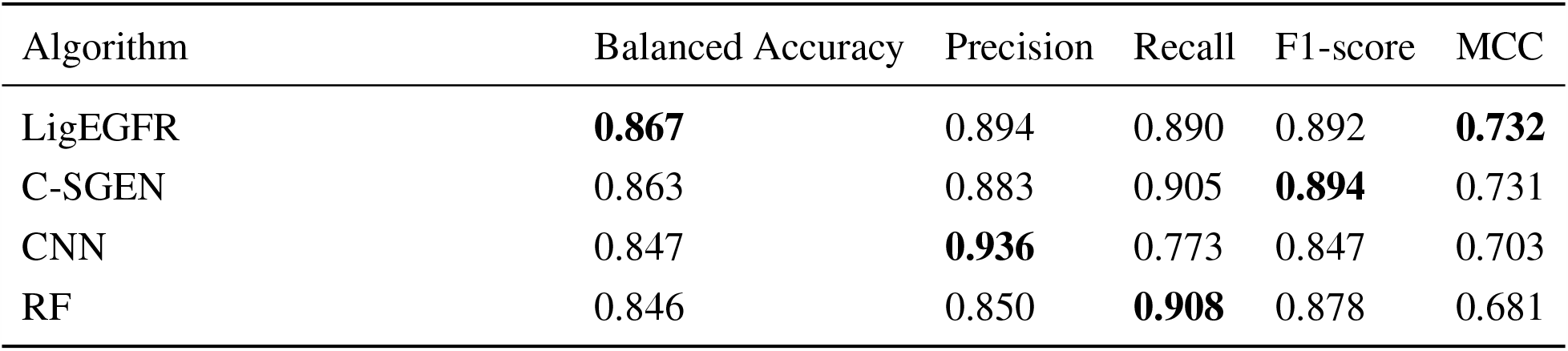
Predictive performance for classification tasks attained by LigEGFR, C-SGEN, CNN, and RF models.

| Algorithm | Balanced Accuracy | Precision | Recall | F1-score | MCC |
| --- | --- | --- | --- | --- | --- |
| LigEGFR | **0.867** | 0.894 | 0.890 | 0.892 | **0.732** |
| C-SGEN | 0.863 | 0.883 | 0.905 | **0.894** | 0.731 |
| CNN | 0.847 | **0.936** | 0.773 | 0.847 | 0.703 |
| RF | 0.846 | 0.850 | **0.908** | 0.878 | 0.681 |

**Figure 8:**
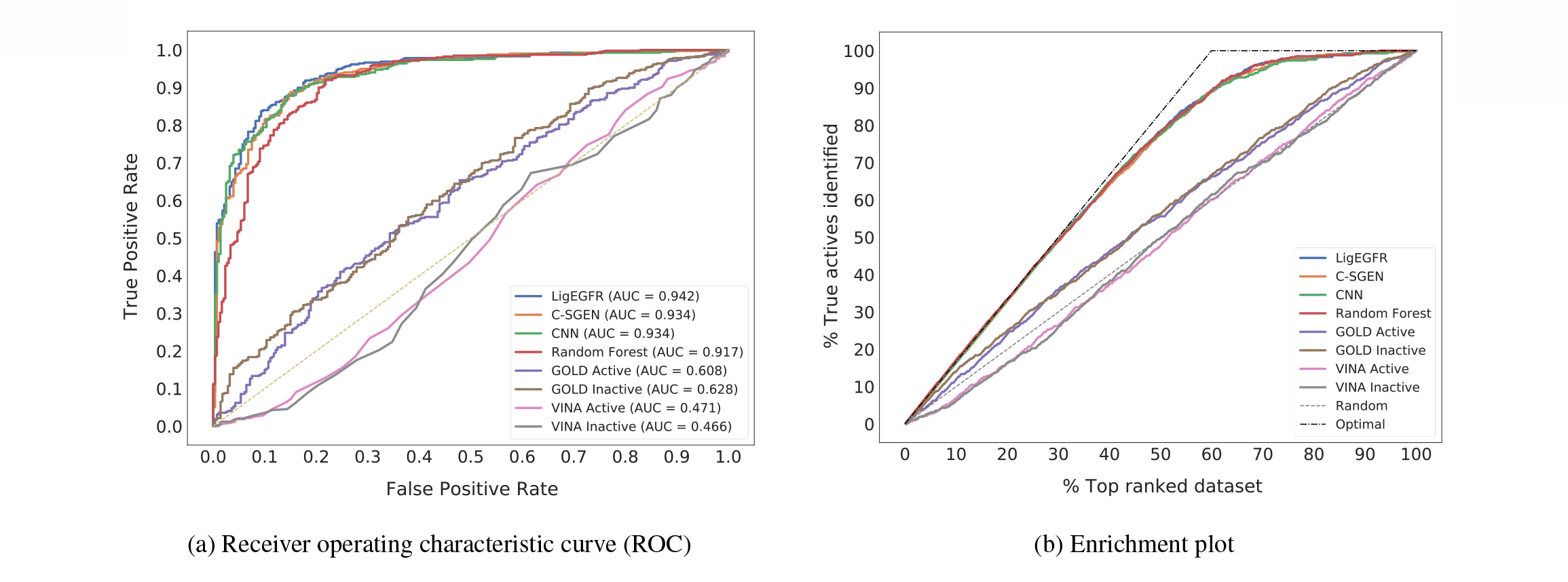
Line plots show (a) the receiver operating characteristic curve (ROC) and (b) the enrichment. For the molecular docking approaches, CCDC GOLD and Autodock Vina, were computed with the active and inactive conformations of human EGFR.

When comparing baselines between the two techniques, machine learning methods have higher performance than the molecular docking methods for human EGFR. By focusing on molecular docking, we endeavored to achieve ligands of the test set for finding the best orientation within binding pockets of the active and inactive conformations of the human EGFR tyrosine kinase. In general, human EGFR functions in the cell membrane with active and inactive conformations. The active conformation is the asymmetric dimer state that EGFR changes to trans-autophosphorylation of specific tyrosine residues on the kinase domain. As for the inactive state, EGFR can be an apparent the symmetric dimer or monomer, which occurs before phosphorylation [59]. Nevertheless, molecular docking has been persistently undertaken to isolate hit compounds. Wang *et al*. [55] evaluated the power of docking by ten software packages. The results showed that molecular docking emerged with a weak correlation between docking scores and experimental binding affinities for the PDBbind dataset (version 2014). This failure is because the molecular docking algorithm is conducted from the “lock and key” concept, which does not take account of the real environment. That is to say, it is done without allowing for binding site flexibility and an aqueous solvent [60]. Interestingly, from the Wang *et al*. report, the flexible docking method, referring to ligand-protein flexible docking, does not appreciably enhance the correlation between the rank of binding affinity and docking score. In contrast, the semi-flexible (ligand-flexible) docking method has been recognized to give better correlations than the flexible docking method for both top-scored and best poses. Indeed, the types of docking can assort into rigid docking, semi-flexible docking, and flexible docking [61]. The rigid docking treats protein and ligand as rigid bodies, while semi-flexible and flexible docking methods consider modifications of bond angles, bond lengths, and torsion angles for molecules. However, the docking scoring function, in terms of binding affinity, is least able to distinguish between active and inactive compounds [62].

Cumulatively, the contribution of the LigEGFR phenomenon is in line with our expectations. This newly developed model contains the graphical information of the molecules, which is helpful to predict pIC_50_ prior to hit compound classification. Consequently, our architecture has a better performance for predicting hit compounds among ranked ligands than the existing methods.

## 4 Conclusion

Here we have proposed the LigEGFR, a novel deep learning architecture for pIC_50_prediction of small molecules against human EGFR. We directly incorporated graphical information of molecules into the embedding layer and trained on the C-SGEN algorithm with architecture adaptation. Additionally, the non-hashed and hashed molecular descriptors have been incorporated in the Conv layers and FC layers, respectively to enhance the predictive performance and generalize the model. The LigEGFR model was trained with various hyperparameters to acquire the optimal model. The experimental evaluations show that LigEGFR has the best performance to predict pIC_50_. Also, LigEGFR tends to outperform the best existing methods for hit compound classification, with high values of balanced accuracy and MCC. These results were obtained by comparing baseline machine learning algorithms and molecular docking methods.

Nowadays, research is focused on machine learning development for pIC_50_ with the human EGFR target. However, these existing methods have used small-scale datasets, containing less diversity of molecular scaffolds and functional groups. Furthermore, the traditional virtual screening such as molecular docking is insufficient to predict hit compounds by either ranking or classifying the hit compound. LigEGFR was developed to overcome these drawbacks and incorporated training on a variety of chemical spaces. Moreover, our model exhibits robustness and reliability, confirmed by y-scrambling evaluation and ADAN, respectively.

The approach that we declare here opens a new way of applying the drug discovery process via targeted lung cancer therapy. Additionally, this approach can help to avoid disadvantages in virtual screening, reducing computation time, and preventing complications that arise from the off-target discovery of novel hit compounds. Lastly, Python executable script of LigEGFR and web service are freely available at https://github.com/scads-biochem/LigEGFR and https://ligegfr.vistec.ist, respectively.

## Supporting information

Supplemental Section S1-S8, Table S1-S5, Figure S1-S3

## 5 Supplementary data

Supplementary data are available online at https://academic.oup.com/bib.

## 6 Code availability and implementation

https://github.com/scads-biochem/LigEGFR

## 7 Authors’ contribution

P.V., N.SA., B.B., J.K., N.SU., and S.N. conceived and planned the machine learning experiments. P.V., N.SA., and B.B. carried out the machine learning experiments. P.V., N.SA., B.B., R.L., and T.S. performed a web service and executable script. B.B., T.N., and T.R. described cancer and molecular biology concepts. P.V., N.SA., B.B., N.SU., T.R., and S.N. contributed to the interpretation of the results and discussion. P.V., N.SA., and B.B. took the lead in writing the manuscript with the consultation with N.SU., T.R., and S.N. This project has initial ideas for machine learning architecture by P.V., N.SA., B.B., J.K., and S.N.

## References

[1] Freddie Bray, Jacques Ferlay, Isabelle Soerjomataram, Rebecca L Siegel, Lindsey A Torre, and Ahmedin Jemal. Global cancer statistics 2018: Globocan estimates of incidence and mortality worldwide for 36 cancers in 185 countries. CA: a cancer journal for clinicians, 68(6):394–424, 2018.

[2] Ping Wee and Zhixiang Wang. Epidermal growth factor receptor cell proliferation signaling pathways. Cancers, 9(5):52, 2017.

[3] F Marc Stewart, Hillard M Lazarus, Paul A Levine, Kathy A Stewart, Imad A Tabbara, and Cynthia A Spaulding. High-dose chemotherapy and autologous marrow transplantation for esthesioneuroblastoma and sinonasal undifferentiated carcinoma. American journal of clinical oncology, 12(3):217–221, 1989.

[4] Roy S Herbst, Daniel Morgensztern, and Chris Boshoff. The biology and management of non-small cell lung cancer. Nature, 553(7689):446–454, 2018.

[5] Si-sheng Ou-Yang, Jun-yan Lu, Xiang-qian Kong, Zhong-jie Liang, Cheng Luo, and Hualiang Jiang. Computational drug discovery. Acta Pharmacologica Sinica, 33(9):1131–1140, 2012.

[6] Yu-Chian Chen. Beware of docking! Trends in pharmacological sciences, 36(2):78–95, 2015.

[7] Sunghwan Kim, Jie Chen, Tiejun Cheng, Asta Gindulyte, Jia He, Siqian He, Qingliang Li, Benjamin A Shoemaker, Paul A Thiessen, Bo Yu, et al. Pubchem 2019 update: improved access to chemical data. Nucleic acids research, 47(D1):D1102–D1109, 2019.

[8] David Mendez, Anna Gaulton, A Patrícia Bento, Jon Chambers, Marleen De Veij, Eloy Félix, María Paula Magariños, Juan F Mosquera, Prudence Mutowo, Michał Nowotka, et al. Chembl: towards direct deposition of bioassay data. Nucleic acids research, 47(D1):D930–D940, 2019.

[9] Michael K Gilson, Tiqing Liu, Michael Baitaluk, George Nicola, Linda Hwang, and Jenny Chong. Bindingdb in 2015: a public database for medicinal chemistry, computational chemistry and systems pharmacology. Nucleic acids research, 44(D1):D1045–D1053, 2016.

[10] Markus A Lill. Multi-dimensional qsar in drug discovery. Drug Discovery Today, 12(23-24):1013–1017, 2007.

[11] Marta M Stepniewska-Dziubinska, Piotr Zielenkiewicz, and Pawel Siedlecki. Development and evaluation of a deep learning model for protein–ligand binding affinity prediction. Bioinformatics, 34(21):3666–3674, 2018.

[12] Hakime Öztürk, Arzucan Özgür, and Elif Ozkirimli. Deepdta: deep drug–target binding affinity prediction. Bioinformatics, 34(17):i821–i829, 2018.

[13] José Jiménez, Miha Skalic, Gerard Martinez-Rosell, and Gianni De Fabritiis. K deep: Protein– ligand absolute binding affinity prediction via 3d-convolutional neural networks. Journal of chemical information and modeling, 58(2):287–296, 2018.

[14] Ingoo Lee, Jongsoo Keum, and Hojung Nam. Deepconv-dti: Prediction of drug-target interactions via deep learning with convolution on protein sequences. PLoS computational biology, 15(6):e1007129, 2019.

[15] Izhar Wallach, Michael Dzamba, and Abraham Heifets. Atomnet: a deep convolutional neural network for bioactivity prediction in structure-based drug discovery. arXiv preprint arXiv:1510.02855, 2015.

[16] Jagat Singh Chauhan, Sandeep Kumar Dhanda, Deepak Singla, Subhash M Agarwal, Gajen-dra PS Raghava, Open Source Drug Discovery Consortium, et al. Qsar-based models for designing quinazoline/imidazothiazoles/pyrazolopyrimidines based inhibitors against wild and mutant egfr. PloS one, 9(7):e101079, 2014.

[17] Hongying Du, Zhide Hu, Andrea Bazzoli, and Yang Zhang. Prediction of inhibitory activity of epidermal growth factor receptor inhibitors using grid search-projection pursuit regression method. PLoS One, 6(7):e22367, 2011.

[18] Shehnaz Fatima and Subhash Mohan Agarwal. Exploring structural features of egfr–her2 dual inhibitors as anti-cancer agents using g-qsar approach. Journal of Receptors and Signal Transduction, 39(3):243–252, 2019.

[19] Silvina E Fioressi, Daniel E Bacelo, and Pablo R Duchowicz. Qsar study of human epidermal growth factor receptor (egfr) inhibitors: conformation-independent models. Medicinal Chemistry Research, 28(11):2079–2087, 2019.

[20] Saw Simeon, Dino Montanari, and Matthew Paul Gleeson. Investigation of factors affecting the performance of in silico volume distribution qsar models for human, rat, mouse, dog & monkey. Molecular informatics, 38(10):1900059, 2019.

[21] Aparna Vema, Sunil K Panigrahi, G Rambabu, B Gopalakrishnan, JARP Sarma, and Gautam R Desiraju. Design of egfr kinase inhibitors: a ligand-based approach and its confirmation with structure-based studies. Bioorganic & medicinal chemistry, 11(21):4643–4653, 2003.

[22] Garima Verma, Mohemmed Faraz Khan, Wasim Akhtar, Mohammad Mumtaz Alam, Mymoona Akhter, Ozair Alam, Syed Misbahul Hasan, and Mohammad Shaquiquzzaman. Pharmacophore modeling, 3d-qsar, docking and adme prediction of quinazoline based egfr inhibitors. Arabian Journal of Chemistry, 12(8):4815–4839, 2019.

[23] Xiaofeng Wang, Zhen Li, Mingjian Jiang, Shuang Wang, Shugang Zhang, and Zhiqiang Wei. Molecule property prediction based on spatial graph embedding. Journal of chemical information and modeling, 59(9):3817–3828, 2019.

[24] Adam Paszke, Sam Gross, Francisco Massa, Adam Lerer, James Bradbury, Gregory Chanan, Trevor Killeen, Zeming Lin, Natalia Gimelshein, Luca Antiga, et al. Pytorch: An imperative style, high-performance deep learning library. In Advances in neural information processing systems, pages 8026–8037, 2019.

[25] Bharath Ramsundar, Peter Eastman, Patrick Walters, Vijay Pande, Karl Leswing, and Zhenqin Wu. Deep Learning for the Life Sciences. O’Reilly Media, 2019.

[26] Reaxys. Reaxys medicinal chemistry. https://www.reaxys.com.

[27] Wes McKinney et al. Data structures for statistical computing in python. In Proceedings of the 9th Python in Science Conference, volume 445, pages 51–56. Austin, TX, 2010.

[28] Stéfan van der Walt, S Chris Colbert, and Gael Varoquaux. The numpy array: a structure for efficient numerical computation. Computing in science & engineering, 13(2):22–30, 2011.

[29] Pauli Virtanen, Ralf Gommers, Travis E Oliphant, Matt Haberland, Tyler Reddy, David Cournapeau, Evgeni Burovski, Pearu Peterson, Warren Weckesser, Jonathan Bright, et al. Scipy 1.0: fundamental algorithms for scientific computing in python. Nature methods, 17(3):261–272, 2020.

[30] John D Hunter. Matplotlib: A 2d graphics environment. Computing in science & engineering, 9(3):90–95, 2007.

[31] Fabian Pedregosa, Gaël Varoquaux, Alexandre Gramfort, Vincent Michel, Bertrand Thirion, Olivier Grisel, Mathieu Blondel, Peter Prettenhofer, Ron Weiss, Vincent Dubourg, et al. Scikitlearn: Machine learning in python. the Journal of machine Learning research, 12:2825–2830, 2011.

[32] G Landrum. Rdkit: Open-source cheminformatics software. GitHub and SourceForge, 10:3592822, 2016.

[33] Charles Tapley Hoyt, Andrej Konotopez, and Christian Ebeling. Pybel: a computational framework for biological expression language. Bioinformatics, 34(4):703–704, 2018.

[34] Michael Waskom, Olga Botvinnik, Joel Ostblom, Maoz Gelbart, Saulius Lukauskas, Paul Hobson, David C Gemperline, Tom Augspurger, Yaroslav Halchenko, John B Cole, et al. mwaskom/seaborn: v0. 10.1 (april 2020). Zenodo, 2020.

[35] Egon L Willighagen, John W Mayfield, Jonathan Alvarsson, Arvid Berg, Lars Carlsson, Nina Jeliazkova, Stefan Kuhn, Tomáš Pluskal, Miquel Rojas-Chertó, Ola Spjuth, et al. The chemistry development kit (cdk) v2. 0: atom typing, depiction, molecular formulas, and substructure searching. Journal of cheminformatics, 9(1):33, 2017.

[36] Thomas N Kipf and Max Welling. Semi-supervised classification with graph convolutional networks. arXiv preprint arXiv:1609.02907, 2016.

[37] Christoph Rücker, Gerta Rücker, and Markus Meringer. y-randomization and its variants in qspr/qsar. Journal of chemical information and modeling, 47(6):2345–2357, 2007.

[38] Johann Gasteiger. Handbook of chemoinformatics. Wiley-VCH, 2003.

[39] Tatiana I Netzeva, Andrew P Worth, Tom Aldenberg, Romualdo Benigni, Mark TD Cronin, Paola Gramatica, Joanna S Jaworska, Scott Kahn, Gilles Klopman, Carol A Marchant, et al. Current status of methods for defining the applicability domain of (quantitative) structure-activity relationships: The report and recommendations of ecvam workshop 52. Alternatives to Laboratory Animals, 33(2):155–173, 2005.

[40] Faizan Sahigara, Kamel Mansouri, Davide Ballabio, Andrea Mauri, Viviana Consonni, and Roberto Todeschini. Comparison of different approaches to define the applicability domain of qsar models. Molecules, 17(5):4791–4810, 2012.

[41] Alexander Tropsha. Best practices for qsar model development, validation, and exploitation. Molecular informatics, 29(6-7):476–488, 2010.

[42] Jang-Sik Choi, My Kieu Ha, Tung Xuan Trinh, Tae Hyun Yoon, and Hyung-Gi Byun. Towards a generalized toxicity prediction model for oxide nanomaterials using integrated data from different sources. Scientific reports, 8(1):1–10, 2018.

[43] Nina Nikolova-Jeliazkova and Joanna Jaworska. An approach to determining applicability domains for qsar group contribution models: an analysis of src kowwin. Alternatives to Laboratory Animals, 33(5):461–470, 2005.

[44] Michael M Mysinger, Michael Carchia, John J Irwin, and Brian K Shoichet. Directory of useful decoys, enhanced (dud-e): better ligands and decoys for better benchmarking. Journal of medicinal chemistry, 55(14):6582–6594, 2012.

[45] Nicolas Triballeau, Francine Acher, Isabelle Brabet, Jean-Philippe Pin, and Hugues-Olivier Bertrand. Virtual screening workflow development guided by the “receiver operating characteristic” curve approach. application to high-throughput docking on metabotropic glutamate receptor subtype 4. Journal of medicinal chemistry, 48(7):2534–2547, 2005.

[46] Kathleen Aertgeerts, Robert Skene, Jason Yano, Bi-Ching Sang, Hua Zou, Gyorgy Snell, Andy Jennings, Keiji Iwamoto, Noriyuki Habuka, Aki Hirokawa, et al. Structural analysis of the mechanism of inhibition and allosteric activation of the kinase domain of her2 protein. Journal of Biological Chemistry, 286(21):18756–18765, 2011.

[47] Edgar R Wood, Anne T Truesdale, Octerloney B McDonald, Derek Yuan, Anne Hassell, Scott H Dickerson, Byron Ellis, Christopher Pennisi, Earnest Horne, Karen Lackey, et al. A unique structure for epidermal growth factor receptor bound to gw572016 (lapatinib): relationships among protein conformation, inhibitor off-rate, and receptor activity in tumor cells. Cancer research, 64(18):6652–6659, 2004.

[48] Andrew Leaver-Fay, Michael Tyka, Steven M Lewis, Oliver F Lange, James Thompson, Ron Jacak, Kristian W Kaufman, P Douglas Renfrew, Colin A Smith, Will Sheffler, et al. Rosetta3: an object-oriented software suite for the simulation and design of macromolecules. In Methods in enzymology, volume 487, pages 545–574. Elsevier, 2011.

[49] Gareth Jones, Peter Willett, Robert C Glen, Andrew R Leach, and Robin Taylor. Development and validation of a genetic algorithm for flexible docking. Journal of molecular biology, 267(3):727–748, 1997.

[50] Oliver Korb, Thomas Stutzle, and Thomas E Exner. Empirical scoring functions for advanced proteinligand docking with plants. Journal of chemical information and modeling, 49(1):84– 96, 2009.

[51] Oleg Trott and Arthur J Olson. Autodock vina: improving the speed and accuracy of docking with a new scoring function, efficient optimization, and multithreading. Journal of computational chemistry, 31(2):455–461, 2010.

[52] Alexander Golbraikh, Min Shen, Zhiyan Xiao, Yun-De Xiao, Kuo-Hsiung Lee, and Alexander Tropsha. Rational selection of training and test sets for the development of validated qsar models. Journal of computer-aided molecular design, 17(2-4):241–253, 2003.

[53] Vladimir Svetnik, Andy Liaw, Christopher Tong, J Christopher Culberson, Robert P Sheridan, and Bradley P Feuston. Random forest: a classification and regression tool for compound classification and qsar modeling. Journal of chemical information and computer sciences, 43(6):1947–1958, 2003.

[54] Svante Wold, Michael Sjöström, and Lennart Eriksson. Pls-regression: a basic tool of chemometrics. Chemometrics and intelligent laboratory systems, 58(2):109–130, 2001.

[55] Zhe Wang, Huiyong Sun, Xiaojun Yao, Dan Li, Lei Xu, Youyong Li, Sheng Tian, and Tingjun Hou. Comprehensive evaluation of ten docking programs on a diverse set of protein–ligand complexes: the prediction accuracy of sampling power and scoring power. Physical Chemistry Chemical Physics, 18(18):12964–12975, 2016.

[56] Dariusz Plewczynski, Michał ŁaŹniewski, Rafał Augustyniak, and Krzysztof Ginalski. Can we trust docking results? evaluation of seven commonly used programs on pdbbind database. Journal of computational chemistry, 32(4):742–755, 2011.

[57] Selcuk Korkmaz. Deep learning-based imbalanced data classification for drug discovery. Journal of Chemical Information and Modeling, 60(9):4180–4190, 2020.

[58] Davide Chicco and Giuseppe Jurman. The advantages of the matthews correlation coefficient (mcc) over f1 score and accuracy in binary classification evaluation. BMC genomics, 21(1):6, 2020.

[59] Ruth Nussinov, Hyunbum Jang, Chung-Jung Tsai, and Feixiong Cheng. Precision medicine and driver mutations: Computational methods, functional assays and conformational principles for interpreting cancer drivers. PLoS computational biology, 15(3):e1006658, 2019.

[60] Pedro J Ballester, Adrian Schreyer, and Tom L Blundell. Does a more precise chemical description of protein–ligand complexes lead to more accurate prediction of binding affinity? Journal of chemical information and modeling, 54(3):944–955, 2014.

[61] Francesca Tessaro and Leonardo Scapozza. How ‘protein-docking’translates into the new emerging field of docking small molecules to nucleic acids? Molecules, 25(12):2749, 2020.

[62] J. Cole, E. Davis, G. Jones, and C.R. Sage. 3.12 - molecular docking—a solved problem? In Samuel Chackalamannil, David Rotella, and Simon E. Ward, editors, Comprehensive Medicinal Chemistry III, pages 297 – 318. Elsevier, Oxford, 2017.

