## Supplemental Section S1-S8, Table S1-S5, Figure S1-S3 for "LigEGFR: Spatial graph embedding and molecular descriptors assisted bioactivity prediction of ligand molecules for epidermal growth factor receptor on a cell line-based dataset"

### S1 Web service construction

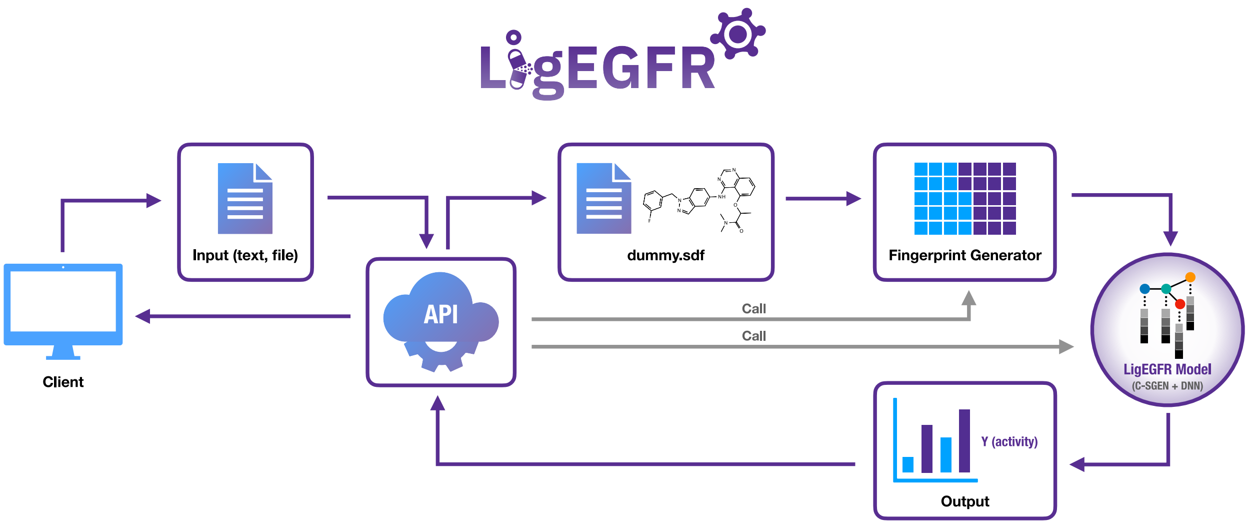


**Figure S1:** The pipeline of the LigEGFR web service contains the LigEGFR architecture for automatic $pIC_{50}$ prediction of small molecules against human EGFR tyrosine kinase.

We constructed the LigEGFR web service for the straightforward prediction of the $pIC_{50}$ and molecular properties of ligands in compliance with the RO5 criterion. This system allows users to generate the molecular structure input in SDF format, preventing the input of unclear chemical structures such as canonical SMILES formats. This website will convert molecular structures in SDF format to isomeric SMILES then calculate fingerprints, and subsequently predict $pIC_{50}$ values by the LigEGFR model. The Vue.js was used for the front-end framework, which has the Netlify platform as a service (PaaS). The Flask platform was used for the back-end framework. The LigEGFR website also provides physicochemical information of the ligand, which is calculated by RDKit. The pipeline of the web construction is shown S1.

**Table S1:** Description of 16 molecular fingerprints from the CDK, PyBEL, and RDKit packages employed in this study.

| Package | Fingerprint | Number of descriptors | Data structure | Description |
|---|---|---|---|---|
| CDK | AtomPair | 2,048 | Hashed | Presence of atomic pairs at specific topological distance |
| | Circular | 2,048 | Hashed | Presence of substructures by considering on neighboring atoms |
| | Daylight | 2,048 | Fragmentation Hashed | Presence of atom types, neighboring atoms, atom groups, and bonds which have lengths from 2-31 bonds |
| | Estate | 79 | Substructure | Presence of substructures which affects the charge of the molecule |
| | Extended | 2,048 | Fragmentation Hashed | Resembles a Daylight fingerprint with additional ring structure |
| | Graph | 2,048 | Fragmentation Hashed | Resembles a Daylight fingerprint but it does not include bond orders |
| | Hybridization | 2,048 | Fragmentation Hashed | Resembles a Daylight fingerprint but it does not consider an aromatic ring |
| | Klekota-Roth | 4,860 | Substructure Decision Tree | Presence of substructures which affect the biological activity |
| | LINGO | 2,048 | Fragmentation Regression | Presence of substructures in SMARTS format which is divided into 4 characters for each substructure |
| | PubChem | 881 | Substructure | Presence of PubChem substructures such as rings, atomic pairs, neighboring atoms, simple & advanced SMARTS formats |
| PyBEL | FP2 | 2,048 | Fragmentation Hashed | Resembles a Daylight fingerprint but it also considers the length of the molecule from 1 to 7 |
| | FP3 | 2,048 | Substructure Hashed | Presence of 204 selected substructures in SMARTS format |
| | FP4 | 2,048 | Substructure Hashed | Presence of 316 selected substructures in SMARTS format |
| RDKit | Avalon | 2,048 | Hashed | Presence of the number of hydrogens, rings, bonds, bond angles, similar to Daylight fingerprint |
| | MACCS | 166 | Substructure | Presence of substructures in SMARTS format |
| | RDKit | 2,048 | Hashed | Presence of RDKit substructures such as atom types, rings, ring sizes, and aromatic rings |

**Table S2:** The details of atomic features in the C-SGEN algorithm.

| Feature | Description | Size | Function |
|---|---|---|---|
| Atom | C, N, O, S, F, Si, P, Cl, Br, Mg, Na, Ca, Fe, As, Al, I, B, V, K, Tl, Yb, Sb, Sn, Ag, Pd, Co, Se, Ti, Zn, H, Li, Ge, Cu, Au, Ni, Cd, In, Mn, Zr, Cr, Pt, Hg, Pb, or Unknown | 44 | *GetSymbol()* |
| Degree | number of bonded neighboring atoms | 11 | *GetDegree()* |
| Implicit valence | number of implicit valences | 7 | *GetImplicitValence()* |
| Formal charge | integer formal charge | 1 | *GetFormalCharge()* |
| Radical electrons | number of radical electrons | 1 | *GetNumRadicalElectrons()* |
| Hybridization | $sp$, $sp^2$, $sp^3$, $sp^3d$, or $sp^3d^2$ | 5 | *GetHybridization()* |
| Aromatic | the atom is a part of an aromatic (*True* or *False*) | 1 | *GetIsAromatic()* |
| Neighbor H | number of hydrogen neighbors | 5 | *GetTotalNumHs()* |

### S2 Hyperparameter tuning of the C-SGEN algorithm for the LigEGFR model

We applied various hyperparameters to the LigEGFR model to find the most suitable model for this dataset. Adaptive Moment Estimation (Adam) was used as an optimizer, and a rectified linear unit (ReLU) was used as the activation function. The supplemental Table S3 shows the hyperparameters that were tuned and the best hyperparameter values. Hyperparameters of the baseline algorithms, CNN and RF, were also adjusted to obtain the highest performance. In each hyperparameter setting, the predictive performance was evaluated as an average value over the five different initial weights and biases.

**Table S3:** Definition of the hyperparameters that were used in the C-SGEN algorithm of the LigEGFR model.

| Hyperparameters | Definition | Hyperparameter Settings* | Best Hyperparameters |
|---|---|---|---|
| *BatchSize* | Number of molecules per batch | 8, 16, 32 | 32 |
| *lr* | Adam learning rate | $5 \times 10^{-6}$, $5 \times 10^{-5}$, $5 \times 10^{-4}$, $5 \times 10^{-3}$ | $5 \times 10^{-5}$ |
| *ch_num* | Number of neurons in C-SGEL | 4, 8, 16, 32 | 4 |
| *k* | Number of filters in *conv1d* | 4, 8, 16 | 8 |
| *csgellayer* | Number of C-SGEL | 1, 2, 3, 4, 5, 6 | 4 |

*Note*: Hyperparameter settings were applied by a random search method.

### S3 Establishing the machine learning baseline via the RF algorithm

We developed the RF model by using a recursive feature elimination (RFE) technique for dimensionality reduction. The *Adjusted* $R^2$ ($R^2_{adj}$) was used as an efficiency indicator to prevent the model from overfitting. The calculation of $R^2_{adj}$ can be executed from $R^2$ of the training set and the number of features used as follows.

$$R^2_{adj} = 1 - \frac{(1 - R^2)(N - 1)}{N - m - 1} \tag{1}$$

where $N$ is the number of compounds, and $m$ are the number of features

RFE is a technique that recursively removes the least important features considered from the Gini index. This technique can find crucial features that make the highest $R^2$. The lowest number of features will be returned for use in the model training. The supplemental Table S4 shows the hyperparameters that were optimized and the best hyperparameter setting.

**Table S4:** Definition of the hyperparameters that were used in the RF algorithm for the baseline model.

| Hyperparameters | Definition | Hyperparameter Settings* | Best Hyperparameters |
|---|---|---|---|
| *bootstrap* | Whether bootstrap samples are used when building trees | *True*, *False* | *False* |
| *max_depth* | The maximum depth of each tree | 20, 40, 60, 80, 100, 200, *None* | 40 |
| *max_features* | The number of features to consider for the best split | *auto*, *sqrt* | *sqrt* |
| *min_samples_leaf* | The minimum number of samples required to be at a leaf node | 1, 2, 5, 10, 20, 50 | 1 |
| *min_samples_split* | The minimum number of samples required to split an internal node | 1, 2, 5, 10, 50 | 5 |
| *n_estimators* | The number of trees in the forest | 200, 400, 600, 800, 1,600, 3,200, 6,400 | 400 |

*Note*: Hyperparameter settings were applied by a binary combination method with greedy search.

### S4 Establishing the machine learning baseline via the CNN algorithm

We employed the CNN algorithm as a baseline model. This model was developed by using feature selection (FS) and principal component analysis (PCA) techniques to reduce the dimension of features. For the FS technique, the molecular descriptors with variance lower than 0.001 were not selected, and the accepted correlation coefficient was changed from 0.75 to 0.90. For the PCA technique, the number of components was varied from 50 to 800. Since there were hashed and non-hashed fingerprints, both FS and PCA were applied to each type. The reduced fingerprints were used as inputs to Conv layers and FC layers as shown below:

1. The hashed and non-hashed fingerprints were applied in the Conv layers and FC layer, respectively.
2. The hashed and non-hashed fingerprints were used in the FC layer and Conv layers, respectively.
3. Both hashed and non-hashed fingerprints (all fingerprints) were applied in both FC and Conv layers.

The supplemental Table S5 shows the hyperparameter settings and the best hyperparameter values for CNN model.

**Table S5:** Definition of the hyperparameters that were used in the CNN algorithm for baseline model.

| Hyperparameters | Definition | Hyperparameter Settings* | Best Hyperparameters |
|---|---|---|---|
| *fisrt_conv* | The dimension of the output from 1st Conv layer | 512, 1024 | 512 |
| *second_conv* | The dimension of the output from 2nd Conv layer | 256, 512 | 256 |
| *third_conv* | The dimension of the output from 3rd Conv layer | 128, 256 | 256 |
| *forth_conv* | The dimension of the output from 4th Conv layer | 64, 128 | 64 |
| *dropout_cnn_rate* | Rate in dropout layer after 4th Conv layer | 0.1, 0.3 | 0.1 |
| *loss* | Loss function | *mean_squared_error*, *mean_absolute_error*, *logcosh* | *mean_squared_error* |
| *activation* | Activation function | *relu*, *linear* | *relu* |
| *BatchSize* | Number of molecules per batch | 8, 16 | 16 |
| *early_patience* | Number of epoch before stopping if validation loss is not decreasing | 10, 20 | 10 |
| *first_fcn* | The dimension of the output from 1st Dense layer | 512, 1024 | 1024 |
| *second_fcn* | The dimension of the output from 2nd Dense layer | 256, 512 | 512 |

*Note*: Hyperparameter settings were applied by a binary combination method with greedy search.

### S5 Establishing the machine learning baseline via the original C-SGEN algorithm

We applied the original C-SGEN algorithm for a controlled experiment. The machine learning model was developed from the conventional C-SGEN algorithm using the combination of hashed and non-hashed fingerprints as input for the DNN. The hyperparameters used in this model were similar to the original ones [1]: *BatchSize* = 8, *lr* = $5 \times 10^{-5}$, *ch_num* =16, *k* = 4, *csgellayer* = 3, *lr_decay* = 0.5, *drop_out_p* = 0.5, *decay_interval* = 10. The predictive performance was evaluated as an average value of over the five different initial weights and biases.

### S6 Establishing the molecular docking baseline via CCDC GOLD

For molecular docking, we performed by CCDC GOLD version 5.8.0 [2] with the test set ligands as input. The human EGFR tyrosine kinases in active and inactive conformations were utilized for the receptor. These receptors were prepared by adding hydrogen atoms, deleting unnecessary water molecules, and deleting ligands. The grid center with $10\,\text{Å}$ from the co-crystal ligand was defined for the binding site of docking [Erlotinib for an active conformation (PDB: 1M17) and Lapatinib for an inactive conformation (PDB: 1XKK)]. The ChemScore Kinase template was employed for parameter optimization of the target. This template allows weak CHO interactions to be considered for weak hydrogen bonds, *e.g.* $\mathrm{CH}\cdots\mathrm{O}$ and N-heterocycle, that is suitable for the kinase protein. The Genetic algorithm (GA) was applied in searching for the best ligand orientation in 150 runs for all ligands of the test set. The no-early termination was considered for virtual screening. The Piecewise Linear Potential (ChemPLP) [3] was dedicated as an empirical fitness scoring function that can be calculated as follows.

$$\mathrm{Fitness}_{\mathrm{ChemPLP}} = \mathrm{Fitness}_{\mathrm{PLP}} - (\mathrm{f}_{\text{chem-hb}} + \mathrm{f}_{\text{chem-cho}} + \mathrm{f}_{\text{chem-met}}) \tag{2}$$

$$\mathrm{Fitness}_{\mathrm{PLP}} = \begin{cases} -(\mathrm{W}_{\mathrm{PLP}}\mathrm{f}_{\mathrm{PLP}} + \mathrm{W}_{\text{lig-clash}}\mathrm{f}_{\text{lig-clash}} + \mathrm{W}_{\text{lig-tors}}\mathrm{f}_{\text{lig-tors}} + \mathrm{f}_{\text{chem-cov}} + \\ \mathrm{W}_{\mathrm{prot}}\mathrm{f}_{\text{chem-prot}} + \mathrm{W}_{\mathrm{cons}}\mathrm{f}_{\mathrm{cons}}) \end{cases} \tag{3}$$

where $\mathrm{f}_{\mathrm{PLP}}$ is the fitness score for PLP

$\mathrm{f}_{\text{chem-hb}}$ is the fitness score for hydrogen bond

$\mathrm{f}_{\text{chem-cho}}$ is the fitness score for CHO weak hydrogen bond

$\mathrm{f}_{\text{chem-met}}$ is the fitness score for metal interaction

$\mathrm{f}_{\text{lig-clash}}$ is the fitness score for heavy-atom clash potential

$\mathrm{f}_{\text{lig-tors}}$ is the fitness score for torsion potential

$\mathrm{f}_{\text{chem-cov}}$ is the fitness score for covalent bond

$f_{chem\text{-}prot}$ is the fitness score for flexible side chain of receptor

$f_{cons}$ is the fitness score for constraint contributions

$W_{PLP}$ is the weight of PLP contributions (1.0 by default)

$W_{lig\text{-}clash}$ is the weight of ligand clash potential (1.0 by default)

$W_{lig\text{-}tors}$ is the weight of ligand torsion potential (2.0 by default)

$W_{prot}$ is the weight of the ChemScore protein potential (1.0 by default)

$W_{cons}$ is the weight of constraint contributions (1.0 by default)

All docking tasks were computed by multi-processing via command-line execution.

### S7 Establishing the molecular docking baseline via AutoDock Vina

We performed molecular docking directly with the test set's ligands by using AutoDock Vina 1.1.2 [4]. The AutoDock Vina uses a genetic algorithm with local gradient optimization (Broyden-Fletcher-Goldfarb-Shanno, BFGS) for the searching algorithm. The receptors and ligands were prepared as PDBQT files, which was done by MGLTools/AutoDockTools version 1.5.7 RC [5]. This step was performed for missing atoms, polar hydrogens, charges, and solvation preparations. A grid box for the binding site of docking was defined by 60 × 60 × 60 points in the *x*, *y*, and *z*-axis direction for both EGFR tyrosine kinase with active (PDB: 1M17) and inactive (PDB: 1XKK) conformations. The coordinate of the center for grid box was set to *x* = 22.0137, *y* = 0.2528, and *z* = 52.7940 for EGFR tyrosine kinase in active conformation, and *x* = 17.1816, *y* = 33.9318, and *z* = 38.4276 for the inactive conformation. The binding mode was adjusted to 150 for efficient searching for the binding candidate. The random seed was set to 1234 for both active and inactive conformations to avoid the non-deterministic nature of the search algorithm. The scoring function of AutoDock Vina is determined by a summation of distance-dependent atom pair interactions as follows.

$$E = \sum w_{t_i t_j}(d_{ij}) \tag{4}$$

$d_{ij}$ is the surface distance calculated by $d_{ij} = r_{ij} - R_{t_i} - R_{t_j}$, where $R_t$ is the van der Waals radius of atom type $t$. $w_{t_i t_j}$ is a weighted summation of steric interactions obtained from all atom pairs, hydrophobic interaction between hydrophobic atoms, and hydrogen bonding.

For $w_{t_i t_j}(d_{ij})$, it can be rewritten to an alternative form for their steric interactions by the equation

$$w_{t_i t_j}(d_{ij}) = \begin{cases} w_1 \times Gauss_1(d) + w_2 \times Gauss_2(d) + \\ w_3 \times Repulsion(d) + w_4 \times Hydrophobic(d) + \\ w_5 \times HBond(d) \end{cases} \tag{5}$$

with the scoring function weights: $w_1 = -0.035579$, $w_2 = -0.005156$, $w_3 = 0.840245$, $w_4 = -0.035069$, and $w_5 = -0.587439$, by default.

The steric and addition terms are shown below:

$$Gauss_1(d) = e^{-(d/0.5\,\text{Å})^2} \tag{6}$$

$$Gauss_2(d) = e^{-((d-3\,\text{Å})/2\,\text{Å})^2} \tag{7}$$

$$Repulsion(d) = \begin{cases} d^2, & \text{if } d < 0\,\text{Å} \\ 0, & \text{if } d \geq 0\,\text{Å} \end{cases} \tag{8}$$

$$Hydrophobic(d) = \begin{cases} 1, & \text{if } d < 0.5\,\text{Å} \\ 1.5\,\text{Å} - d, & \text{if } 0.5\,\text{Å} > d < 1.5\,\text{Å} \\ 0, & \text{if } d > 1.5\,\text{Å} \end{cases} \tag{9}$$

$$HBond(d) = \begin{cases} 1, & \text{if } d < -0.7\,\text{Å} \\ \frac{d}{0.7\,\text{Å}}, & \text{if } -0.7\,\text{Å} < d < 0\,\text{Å} \\ 0, & \text{if } d > 0\,\text{Å} \end{cases} \tag{10}$$

### S8 Feature analysis of molecular descriptors

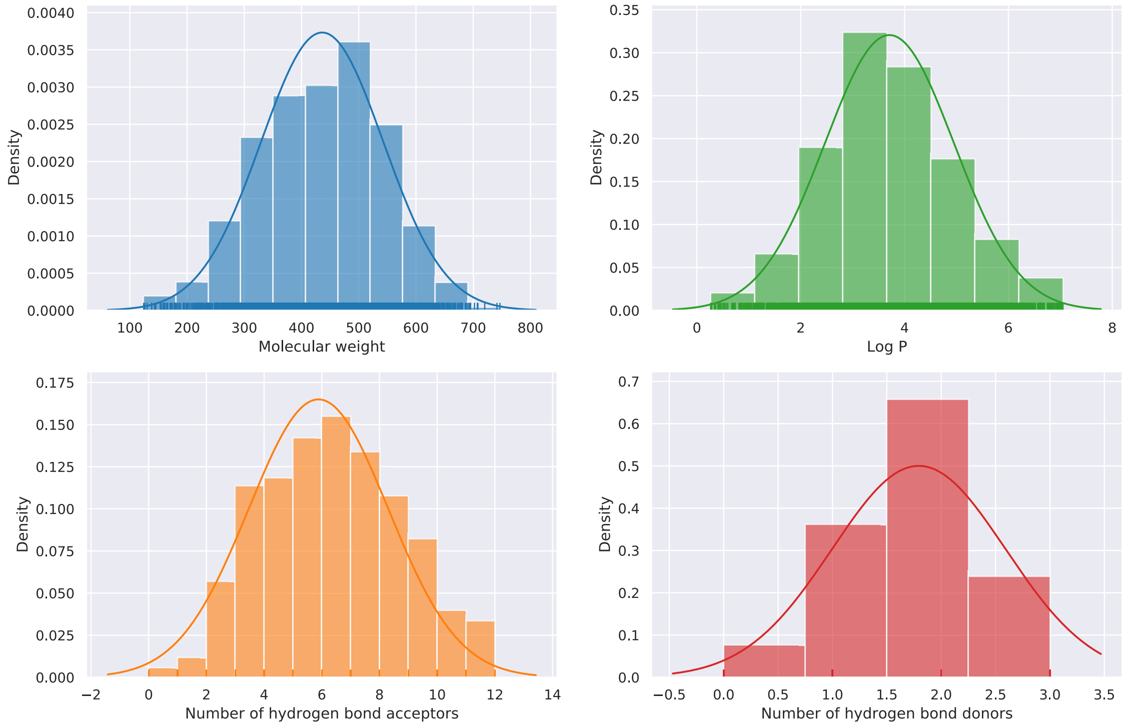


**Figure S2:** Histogram plots of the dataset distributions for molecular weight, $Log$ P, number of hydrogen bond acceptors, and number of hydrogen bond donors. The outliers were removed by using the interquartile range method (IQR).

We retrieved bioactivity data of wild type EGFR from the REAXYS database [6], containing medicinal chemistry information of compounds. The raw data of 37,753 substances were cleaned by selecting only crucial information and removing outliers. As a result, 3,493 substances were considered for machine learning development. The histograms of the density plotted against the RO5 criteria after deleting outliers is shown as Figure S2. The dataset in Gaussian distribution shows that most of the outliers were removed. Furthermore, most of the data are in RO5, with the number of hydrogen bond donors $\leq 5$, the number of hydrogen bond acceptors $\leq 10$, molecular weight $< 500$ daltons, and octanol-water partition coefficient ($Log$ P) $\leq 5$ [7]. Note that the RO5 is a rule of thumb for determining the lead- and drug-like compounds for oral administration that are conserved to the drug's pharmacokinetics, including absorption, distribution, metabolism, and elimination (ADME). The most orally active compounds tend to overemphasize this guideline and have accomplished phase II clinical trial phase [8]. Despite this, the protein kinase inhibitors

(PKIs) from the ChEMBL database may show RO5 violation [9]. Our dataset has contained small molecules with properties falling outside RO5 boundaries; it can predict various physicochemical properties.

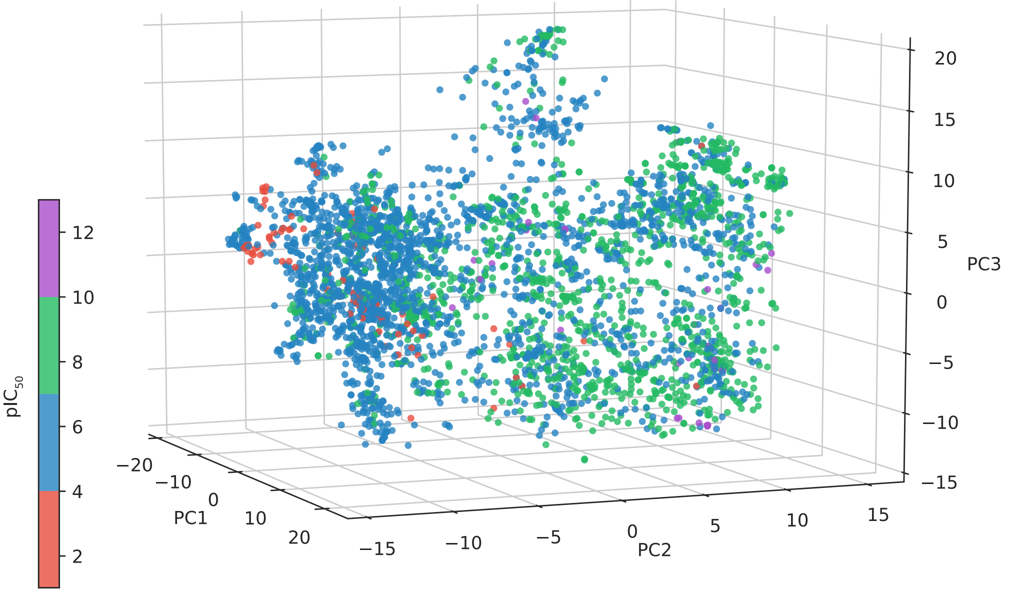


**Figure S3:** The 3D PCA from the molecular descriptors used in LigEGFR. The color of each dot is determined by the $pIC_{50}$ value.

The feature analysis was calculated with 3D principal component space (see Figure S3). Each point color shows the $pIC_{50}$ value. The result indicates that the features have broadly distinguished the group of ligands according to the $pIC_{50}$ values. Hence, they can effectively represent the structure of ligands in the dataset and affect the $pIC_{50}$ prediction.

### References

[1] Xiaofeng Wang, Zhen Li, Mingjian Jiang, Shuang Wang, Shugang Zhang, and Zhiqiang Wei. Molecule property prediction based on spatial graph embedding. *Journal of chemical information and modeling*, 59(9):3817–3828, 2019.

[2] Gareth Jones, Peter Willett, Robert C Glen, Andrew R Leach, and Robin Taylor. Development and validation of a genetic algorithm for flexible docking. *Journal of molecular biology*, 267(3):727–748, 1997.

[3] Oliver Korb, Thomas Stutzle, and Thomas E Exner. Empirical scoring functions for advanced protein- ligand docking with plants. *Journal of chemical information and modeling*, 49(1):84–96, 2009.

[4] Oleg Trott and Arthur J Olson. Autodock vina: improving the speed and accuracy of docking with a new scoring function, efficient optimization, and multithreading. *Journal of computational chemistry*, 31(2):455–461, 2010.

[5] Garrett M Morris, Ruth Huey, William Lindstrom, Michel F Sanner, Richard K Belew, David S Goodsell, and Arthur J Olson. Autodock4 and autodocktools4: Automated docking with selective receptor flexibility. *Journal of computational chemistry*, 30(16):2785–2791, 2009.

[6] Reaxys. Reaxys medicinal chemistry. `https://www.reaxys.com`.

[7] Christopher A Lipinski, Franco Lombardo, Beryl W Dominy, and Paul J Feeney. Experimental and computational approaches to estimate solubility and permeability in drug discovery and development settings. *Advanced drug delivery reviews*, 23(1-3):3–25, 1997.

[8] Christopher A Lipinski. Lead-and drug-like compounds: the rule-of-five revolution. *Drug Discovery Today: Technologies*, 1(4):337–341, 2004.

[9] Colin Bournez, Fabrice Carles, Gautier Peyrat, Samia Aci-Sèche, Stéphane Bourg, Christophe Meyer, and Pascal Bonnet. Comparative assessment of protein kinase inhibitors in public databases and in pkidb. *Molecules*, 25(14):3226, 2020.
